# Pig vocalizations contain shared acoustic structure for humans and machines, but limited correspondence with presumed affective valence

**DOI:** 10.64898/2026.07.06.736900

**Authors:** Wim Gorssen, Ben Sleurs, Carmen Winters

## Abstract

Pig vocalizations are increasingly used as indicators of affective state and welfare. This view is supported by above-chance human valence judgments and striking machine learning classification performance in prior studies. Yet, in both approaches the affective distinction is already built into the test. Algorithms are trained and evaluated using presumed valence labels assigned to recording contexts, while human listeners are asked directly to rate valence. We therefore asked what structure emerges when listeners are given no predefined categories, and how that structure relates to explicit human judgments and neural network representations. In a free-classification task, 224 participants labeled and sorted 2,192 pig vocalizations. A second group of 159 participants then classified vocalizations using predefined context categories and rated their valence. Humans and neural networks recovered corresponding organization, confirming that pig vocalizations contain reproducible acoustic information. Yet, affect was not the dominant spontaneous interpretation, with only 26.3% of participant-generated labels referring to feelings or emotions. Explicit valence judgments were systematic and agreed moderately with presumed valence, with 60.1% directional agreement. However, this correspondence disappeared when highly aversive situations were excluded, although neural networks still classified presumed valence with 92.0% accuracy after removing these contexts. In a matched permutation analysis, however, neural networks with arbitrary valence reassignments on the context level averaged 79% accuracy, with 5 of 100 exceeding the original grouping. Pig vocalizations therefore contain rich acoustic information that both humans and algorithms can recover, but successful recovery does not by itself establish what that information means for the animal. Progress in vocal indicators of pig affect and welfare will require moving beyond classification performance toward independent validation of the biological meaning assigned to acoustic differences.

**Significance Statement:** Animal vocalizations are increasingly used to infer affective state and welfare, yet presumed valence is often assigned at the level of the recording context and then applied to individual vocalizations produced within that context. We show that humans and machine-learning models recover similar acoustic organization in pig vocalizations, especially for highly aversive situations, but that this organization only partially corresponds to context-derived valence. More fundamentally, successful classification of presumed affective state is not equivalent to demonstrating that vocalizations provide a valid readout of the animal’s affective state. Their use as welfare indicators therefore requires independent validation against animal-derived evidence of what animals seek or avoid and consequences for their health, rather than inferred from human-assigned emotional categories.

## Introduction

Animal vocalizations are increasingly investigated as potential indicators of animal affective state and welfare (Villain & Briefer, 2026; Nicolaisen et al., 2025; Whitham et al., 2024; Briefer et al., 2012). An influential perspective is that vocal expressions are linked to the producer’s affective state and can therefore provide indicators or potentially real-time measures to study affective state (Villain & Briefer, 2026). In much pig vocalization research, this hypothesis has been tested by assigning presumed affective valence a priori at the recording-context level, such that individual vocalizations inherit the valence assigned to that context (Briefer et al., 2019; Friel et al., 2019; Tallet et al., 2013). Here, valence broadly refers to the positive or negative quality of subjective experience. Affective valence has been represented categorically in some pig vocalization studies (Nicolaisen et al., 2025), but more commonly within a positive-to-negative valence framework (See Table S1; Briefer et al., 2019, Friel et al., 2019; Maigrot et al., 2018).

Machine-learning approaches have further strengthened the impression that affective state can be recovered directly from pig vocalizations. Models trained on context-level valence or stress-related labels have achieved moderate to high classification performance, including 91.5% accuracy for presumed positive versus negative contexts (Briefer et al., 2022a), 83.9% balanced accuracy in a multispecies valence classifier (Lefèvre et al., 2025), and 60.9% balanced accuracy for experimentally imposed distress conditions in previously unseen pigs (Nääs et al., 2026). Human listeners have likewise rated presumed valence above-chance levels, but under tasks that explicitly directed them toward particular affective or contextual interpretations. Tallet et al. (2010) asked listeners to classify pig vocalizations within four production contexts and rate their affective valence, whereas Greenall et al. (2022) asked listeners to discriminate vocalizations originating from presumed positive or negative conditions.

Recent syntheses have interpreted this literature as providing strong construct validity for vocal indicators of affective valence and supporting their use in welfare assessment (Villain & Briefer, 2026). However, the evidence is considerably stronger that vocalizations differ between experimental conditions than that these differences warrant inference about the affective valence of the individual animal. Moving from one claim to the other requires five additional assumptions about what the contexts represent, what drives the acoustic differences, and how well these relationships generalize. We therefore evaluated these assumptions in an inferential audit of 16 studies (Table 1; Table S1). None of the studies simultaneously established the animal’s momentary valence and demonstrated that the acoustic effect was specific to valence.

**Table 1.** Inferential audit of evidence linking pig vocalizations to affective state. Studies were evaluated against five auxiliary assumptions required to interpret vocal variation as evidence of affective state: **A1, construct realization**, whether the experimental manipulation or observational context produced the proposed affective state; **A2, construct comparability**, whether states elicited by different contexts can be represented meaningfully on a common affective dimension; **A3, construct specificity**, whether the vocal effect can be attributed specifically to the proposed affective construct rather than correlated processes such as arousal, pain, motivation, activity, social context, or call function; **A4, measurement correspondence**, whether the observed vocal relationship corresponds to variation in the state of the vocalizing individual rather than only to group- or context-level differences; and **A5, transportability**, whether the relationship persists across animals, call types, eliciting contexts, developmental stages, populations, or recording settings. The final column reports the **highest supported claim** without requiring unresolved affective assumptions. **C0** denotes evidence that vocalizations differ among experimentally or observationally defined conditions, ‘-‘ is filled in if the study did not study affect in specific (e.g., stress or arousal); higher claim levels concern association with affective change, valence specificity, cross-context robustness, and prediction of affective state in novel contexts.

| Study | A1<br>Construct<br>realization | A2<br>Construct<br>comparability | A3<br>Construct<br>specificity | A4<br>Measurement<br>correspondence | A5<br>Transportability | C<br>Highest<br>supported<br>claim |
| --- | --- | --- | --- | --- | --- | --- |
| Schrader, 1998 | ● Direct* | ● Inconclusive | ● Inconclusive | ● Inconclusive | ● Contrasted | - |
| Imfeld-Mueller, 2011 | ● Inconclusive | ● Inconclusive | ● Inconclusive | ● Direct<sup>1</sup> | ● Not tested | C0 |
| Reimert, 2013 | ● Inconclusive | ● Inconclusive | ● Inconclusive | ● Not tested | ● Not tested | C0 |
| Tallet, 2013 | ● Inconclusive | ● Inconclusive | ● Not tested | ● Not tested | ● Direct<br>● Not tested (broader transportability) | C0 |
| Linhart, 2015 | ● Inconclusive | ● Inconclusive | ● Inconclusive | ● Direct<sup>1</sup> | ● Contrasted (cross call-type)<br>● Not tested (cross-context) | - |
| Leliveld, 2016 | ● Inconclusive | ● Inconclusive | ● Contrasted | ● Inconclusive | ● Contrasted | C0 |
| Maigrot, 2018 | ● External | ● Inconclusive | ● Inconclusive | ● Direct<sup>1</sup> | ● Contrasted | C0 |
| Briefer, 2019 | ● Inconclusive | ● External | ● Inconclusive | ● Direct<sup>1</sup> | ● Inconclusive | C0 |
| Friel, 2019 | ● Inconclusive | ● Inconclusive | ● Inconclusive | ● Direct<sup>1</sup> | ● Inconclusive | C0 |
| Villain, 2020a | ● Inconclusive | ● Inconclusive | ● Inconclusive | ● Direct<sup>1</sup> | ● Inconclusive | C0 |
| Villain, 2020b | ● Inconclusive | ● Inconclusive | ● Inconclusive | ● Direct<sup>1</sup> | ● Contrasted | C0 |
| Briefer, 2022 | ● External | ● Inconclusive | ● Inconclusive | ● Inconclusive | ● Direct*(within dataset)<br>● Not tested (broader transportability) | C0 |
| Lefèvre, 2025 | ● External | ● Inconclusive | ● Inconclusive | ● Inconclusive | ● Direct* | C0 |
| Nicolaisen, 2025 | ● Inconclusive | ● Inconclusive | ● Inconclusive | ● Not tested | ● Inconclusive | C0 |
| Nääs, 2026 | ● Inconclusive | ● Inconclusive | ● Inconclusive | ● Direct<sup>1</sup> | ● Direct* (moderate cross-animal)<br>● Contrasted (age-invariant mapping)<br>● Not tested (cross-context) | C0 |
| Valfré, 2026 | ● Inconclusive | ● Inconclusive | ● Inconclusive | ● Inconclusive | ● Not tested | C0 |

\*Statuses describe the evidence available for each auxiliary assumption rather than overall study quality. **Direct** indicates that the assumption was directly addressed within the study; **External** indicates that support was imported primarily from previous literature or theory; **Inconclusive** indicates that the available evidence did not resolve the assumption; **Contrasted** indicates that direct evidence showed that the assumed relationship was not invariant across the tested conditions; and **Not tested** indicates that the assumption was required for the relevant inference but was not meaningfully evaluated.
<sup>1</sup>Directly addresses within-individual/context-linked acoustic variation, not independently measured within-individual valence.

More fundamentally, assigning a single valence to a context does not establish that the presumed valence was experienced by every individual animal, or that it applied uniformly throughout the evaluated recording period. Context labels therefore do not by themselves establish that changes in vocal acoustics track changes in the individual’s affective state (e.g. Schräder and Todt, 1998). Studies where within-individual, cross-context, or developmental comparisons were available, showed that acoustic relationships were often dependent on context, call type, acoustic parameter, or age (Nääs et al., 2026; Linhart et al., 2015; Illmann et al., 2013). Although Nääs et al. (2026) demonstrated moderate generalization to previously unseen pigs, broader generalization across contexts, populations, and settings remained limited or untested. Thus, current evidence robustly supports condition-related acoustic differences, but not yet a unique, dynamic, and generalizable mapping between vocal acoustics and individual affective valence.

Importantly, this distinction does not depend on denying that non-human animals have affective experiences, nor on assuming that vocalizations contain no affect-relevant information. The narrower question is what evidence warrants identifying affective valence as the construct underlying an observed acoustic relationship (Winters, 2026). The construct validation framework applied by Villain and Briefer (2026) tests whether candidate indicators covary with preferences and avoidance, fitness-relevant situations, pharmacological manipulations, human self-reported states, or other putative affect indicators (Mason & Mendl, 2026). However, these tests support an affective interpretation only insofar as the five auxiliary assumptions linking each observable manipulation or correlate to affective valence are themselves justified (Table 1; Table S1). These assumptions matter because the observations count as evidence for valence only conditional on them. If the same observations are compatible with other biological processes, the data warrant the observable association but not yet the valence-specific interpretation. Convergence among such tests can therefore strengthen evidence for a reproducible relationship without necessarily discriminating affective valence from alternative processes such as motivation, arousal, pain, social context, or communicative function. Validation of an acoustic relationship and validation of its affective interpretation are therefore treated here as related but distinct inferential problems.

This inferential problem is consistent with evidence in humans that mappings between internal states, vocal acoustics, and perceived emotion are probabilistic and context-dependent rather than one-to-one (Barrett, 2025; Gendron et al., 2014). Similar vocal patterns can arise from different underlying processes, while similar affective states may produce different expressions, complicating inference from signal to latent state (Barrett, 2025). Although constructionist perspectives were developed primarily to explain human emotion perception, they have also been proposed as a framework for studying affect in nonhuman animals (Bliss-Moreau, 2017).

The second important limitation of studies in which human listeners rated the valence of pig vocalizations (Tallet et al., 2010; Greenall et al., 2022) is their reliance on predefined response frameworks, including valence rating scales and predefined context categories. Such tasks can show that listeners make systematic affective judgments when asked to do so, but they cannot establish that affective meaning is an inherent property of the vocalization itself. This concern is supported by research on human emotion perception showing that the interpretation of vocal signals can be shaped by the labels and contextual information available to the listener (Gendron et al., 2012, 2014; Lindquist et al., 2006). Moreover, forced-choice designs generally require participants to assign each stimulus to one of the available categories or scales, providing limited opportunity to express uncertainty or to organize stimuli along alternative dimensions. Consequently, forced-choice performance alone cannot distinguish affective information that emerges from the vocalization from affective interpretation shaped by the task.

This study therefore examines how humans organize pig vocalizations when no predefined categories are provided, and how their judgments change when context and valence are introduced as response options (Fig. 1). In Wave 1, participants freely classified pig vocalizations, allowing us to examine the perceptual structure that emerged and the meanings participants assigned to their own categories. In Wave 2, participants classified vocalizations using predefined response frameworks, allowing direct comparison between free and forced-choice task formats. We further tested whether human-derived perceptual structure converged with acoustic structure learned by convolutional neural network models. Together, these analyses allow us to distinguish acoustic structure, perceptual organization, and affective interpretation.

**Figure 1.**
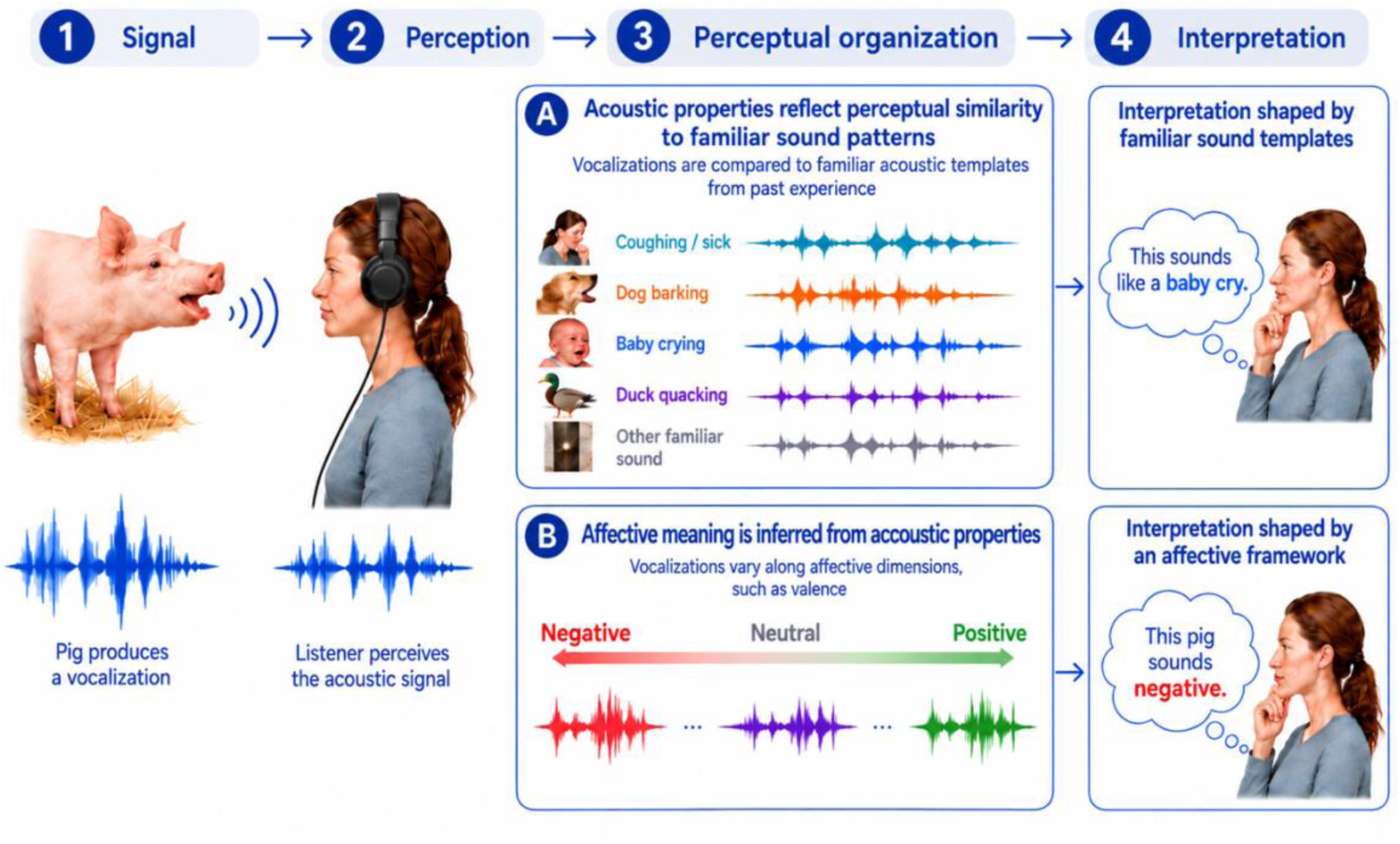
Conceptual framework for human interpretation of pig vocalizations. Human judgments of animal vocalizations can be considered at different levels from acoustic signal production to perception, perceptual organization, and interpretation. One possibility is that acoustic similarity supports broader perceptual grouping (3A), with vocalizations compared to familiar sound patterns or acoustic templates from past experience, which may subsequently shape interpretation. An alternative, non-mutually exclusive possibility is that acoustic properties directly support inference of affective meaning (3B), for example along a valence dimension from negative to positive, within an affective interpretive framework. This framework motivates our comparison of free classification with researcher-defined context classification and valence ratings. Figure generated with assistance from ChatGPT (OpenAI).

## Results

### Human classification revealed reproducible broad perceptual structure across pig vocalization contexts

Across two waves, human participants classified pig vocalizations using either free classification or forced-choice tasks. In a first survey, 224 participants freely sorted 8,960 presentations of 2,192 unique vocalizations into 1,273 unique participant-defined categories. In a second survey, 159 participants classified 3,975 presentations of 1,357 unique vocalizations into predefined context categories and a five-point valence scale.

Human free classification revealed broad but incomplete perceptual structure. Labels were moderately consistent within recording contexts (mean consistency of 45.2%, SD = 17.9%; Fig. S1). Duplicate (castrated piglet) and outlier (dolphin) controls were classified highly consistently (92.9% and 93.8%, respectively), indicating that the task was interpretable and that low consistency for many contexts was not simply due to participant inattention.

Co-classification similarities showed that participants did not recover the original experimental contexts as discrete categories. Castration and restraint showed the highest co-classification similarity (0.88), with 88% of participants placing at least one vocalization from each context in the same self-defined category (Fig. 2 and Fig. S2). Contexts grouped into broad families: high-intensity aversive contexts, especially castration, restraint, fighting, and crushing, grouped most clearly, while structure among the remaining contexts was more graded. Some of this organization also appeared to coincide with age category. For example, several nursing-related contexts occupied a similar region of the ordination. The global co-classification structure was highly similar across expertise groups (Mantel correlation = 0.92–0.94), indicating that pig-related experience had little influence on the broad perceptual organization.

**Figure 2.**
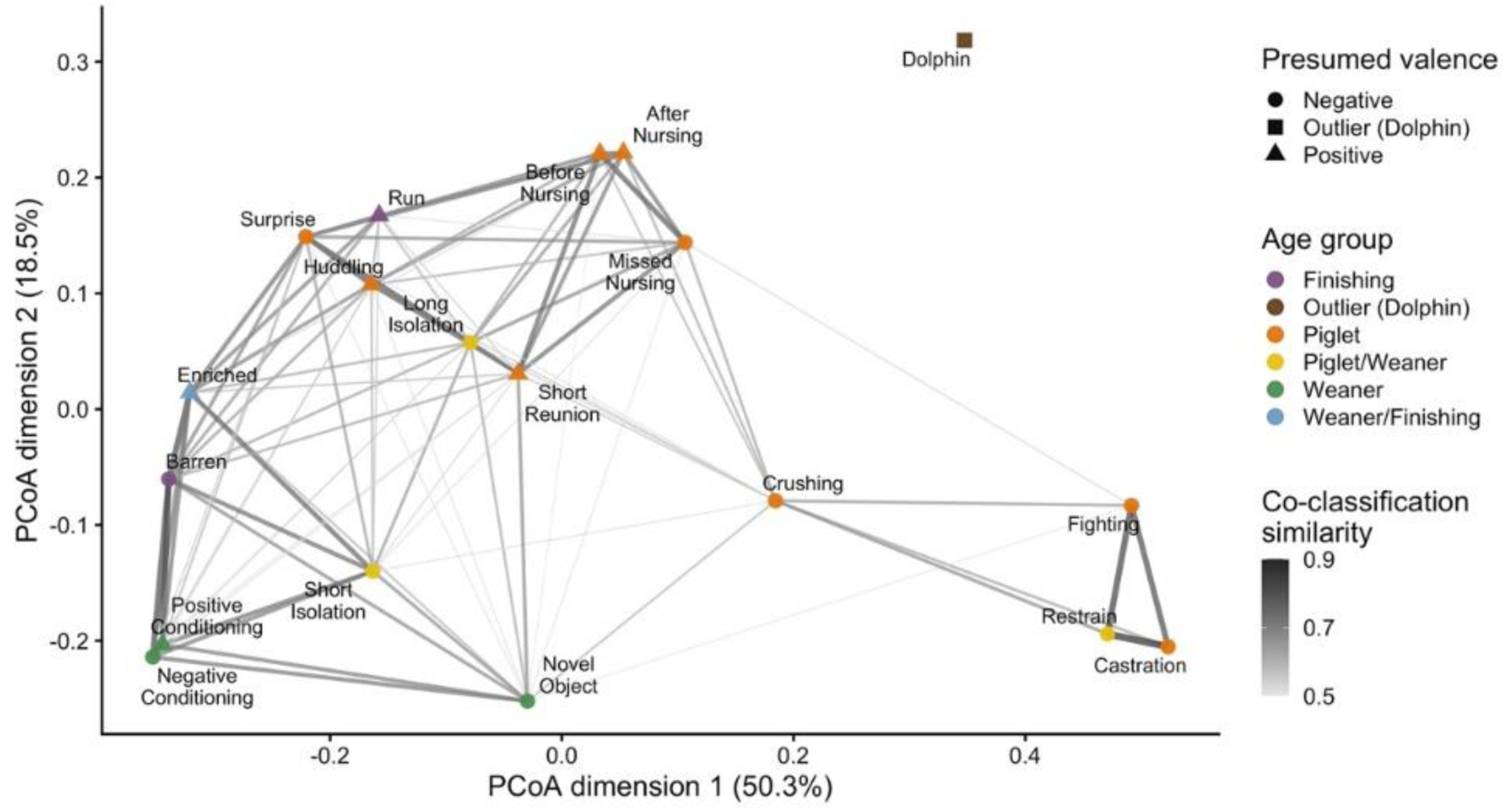
Human perceptual similarity among pig vocalization contexts in Wave 1. Principal coordinates analysis of free-classification similarity among the true recording contexts as described in Briefer et al. (2022a). Points represent contexts, and distances reflect dissimilarity in participant co-classification patterns. Lines connect context pairs with similarity ≥ 0.50, indicating that at least 50% of participants classified sounds from these contexts together, with thicker lines indicating stronger similarity. Point color indicates age group and point shape indicates presumed valence as in Briefer et al. (2022a). Related heatmaps and PCoA plots for different classification scenarios are shown in Figs. S2–S10.

The perceptual structure observed in free classification was also reflected in the forced-choice tasks from Wave 2 for context (Figs. S3-4) and valence (Figs. S5-6). Mantel tests comparing context-to-context distance matrices showed strong, significant correlations (P<0.0001) between free classification and forced-choice context classification (r = 0.82), between free classification and forced-choice valence scoring (r = 0.90), and between forced-choice context classification and valence scoring (r = 0.84). Thus, although participants did not classify vocalizations into discrete recording contexts with high specificity, the three task formats converged on a reproducible broad perceptual organization of pig vocalizations.

### Free-classification labels were mostly descriptive, with sparse valence information driven by negative labels

A language analysis was performed on the participant-defined labels of the free classification task in Wave 1. Participant-defined labels were more often descriptive than explicitly related to feelings or emotions (26.3% of labels). Across free labels, 31.1% were classified as states, 30.6% as animal-related labels (of which 8.5% pig-related), 22.4% as feelings, 15.6% as events, and only 4.6% as emotion labels. Semantic labels often co-occurred within categories, with the highest overlaps between feeling and state (13%; Fig. 3). At the participant level, 81.2% of participants produced at least one state label, 63.4% at least one animal label, 63.4% at least one feeling label, 40.2% at least one event label, and 20.5% at least one emotion label.

**Figure 3.**
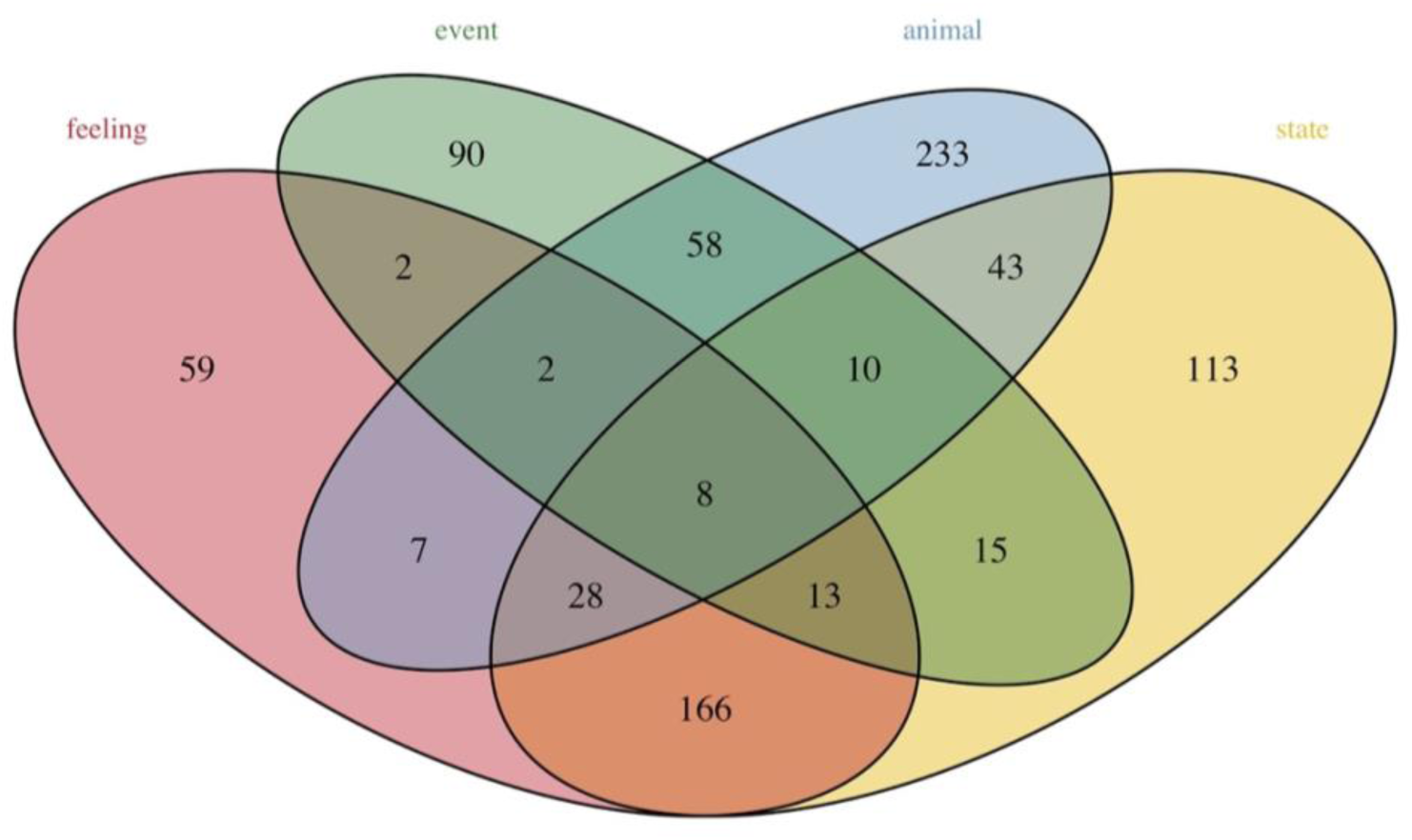
Venn-diagram of overlap in label category from free classification language analysis. A total of 426 of 1,273 unique participant-generated labels (33.5% of total) did not belong to any of these four broad categories.

Professional background affected the type of labels used. Compared with lay participants, pig farmers and veterinarians had lower odds of using feeling labels (pig farmers: OR = 0.42, P < 0.0001; veterinarians: OR = 0.78, P = 0.005), whereas animal scientists had slightly higher odds (OR = 1.19, P = 0.003). In contrast, participants with professional backgrounds in veterinary medicine or animal science were more likely to use event labels (veterinarians: OR = 2.31, P < 0.0001; animal scientists: OR = 2.14, P < 0.0001), and pig farmers were much more likely to use pig-related animal labels (OR = 5.26, P < 0.0001).

The proportion of categories labeled as feeling varied across vocalization contexts, with a mean of 20.6% and ranging from 14.1% (surprise) to 35.9% (castration). Negative feeling labels were most prevalent over contexts (mean = 12.7%; min = 5.1%; max = 34.4%), while positive feeling labels were less prevalent (mean = 5.2%; min = 0.9%; max = 8.9%). Feeling labels were most common and predominantly negative for the contexts castration (35.9% assigned as feeling of which 95.7% as negative), restraint (35.3% assigned as feeling of which 92.4% as negative) and fighting (29.7% assigned as feeling of which 86.5% as negative) (Table S2). Table S2 also shows that none of the contexts had vocalizations that were predominantly freely labeled as positive by human listeners.

We then asked whether spontaneous valence labels in Wave 1 aligned with presumed valence by Briefer et al. (2022a). When positive or negative free labels were used, 66.7% matched the presumed direction, but this signal was asymmetric. Negative labels were more frequent and more precise than positive labels (74.8% versus 45.4%), and conditional recall was higher for presumed negative than presumed positive sounds (78.3% versus 40.6%). After removing high-intensity aversive contexts, directional alignment dropped to chance-like levels, with overall precision of 49.6%. Thus, free classification contained a sparse valence signal that was driven mainly by negative labels from high-intensity aversive contexts.

### Valence judgments converged across free- and forced-choice tasks

Feeling- and emotion-related labels from Wave 1 were converted to a signed valence proxy, with positive labels coded as 1, negative labels as −1, and unclear or non-valence categories coded as 0. This proxy was positively associated with explicit Wave 2 valence ratings for the same sounds (Spearman’s ρ = 0.38, P < 0.0001; Fig. 4A). Importantly, the association strengthened as analyses were restricted to sounds rated by more participants in both waves, reaching ρ = 0.72 for sounds rated by at least four listeners per wave (P < 0.0001; Fig. 4D). Thus, despite the different task formats and participant samples, sounds that elicited more positive or negative spontaneous labels in Wave 1 received correspondingly more positive or negative explicit ratings in Wave 2.

**Figure 4.**
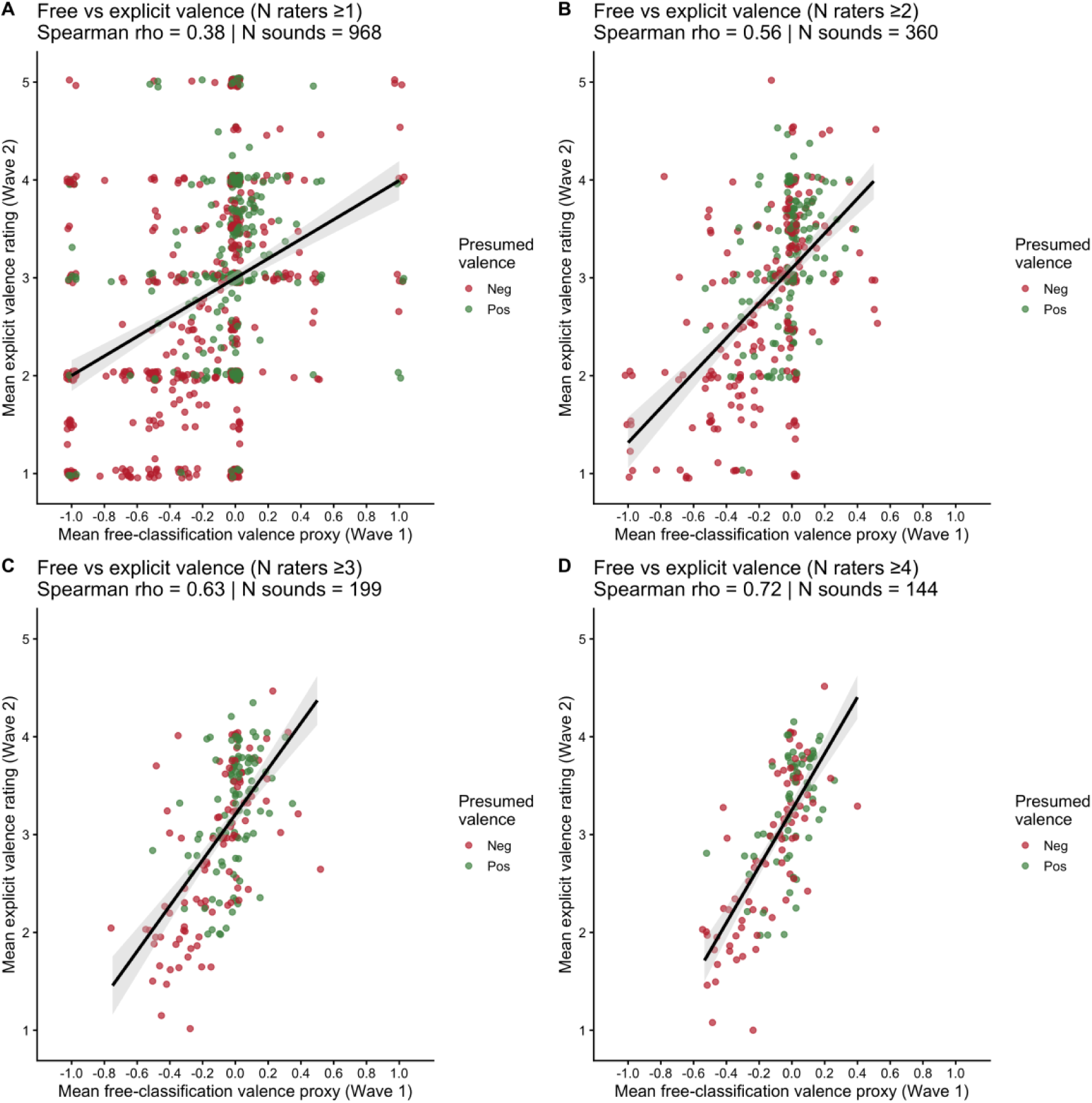
Valence judgments converge across free-classification and explicit valence-rating tasks. Relationship between a valence proxy derived from Wave 1 free-classification labels and explicit Wave 2 valence ratings for the same vocalizations. Feeling- and emotion-related free labels were converted to a signed proxy score, with positive labels coded as 1, negative labels as −1, and mixed labels or labels without feeling or emotion labels as 0. Explicit Wave 2 valence ratings were scored on a five-point Likert scale from very negative to very positive. Each point represents the average values for one sound over participants. Points were jittered to show overlapping points and colored by the presumed valence as defined by Briefer et al. (2022a). Panels show increasingly strict inclusion thresholds based on the minimum number of raters (N raters) per sound in both waves: at least one rater per wave (A), at least two raters per wave (B), at least three raters per wave (C), and at least four raters per wave (D). Blue lines show linear fits with 95% confidence intervals. Spearman’s ρ (rho) and the number of sounds (N sounds) included are shown within each panel.

After excluding vocalizations from castration, restraint, fighting, and crushing, the association remained strong for sounds rated by at least four listeners per wave (ρ = 0.61, P < 0.0001; Fig. S11). These results demonstrate substantial cross-task consistency in human valence judgments of individual pig vocalizations, even when agreement with the presumed valence categories assigned to the recording contexts was weak.

A parallel analysis of uncertainty showed a similar cross-task pattern: sounds that elicited more explicit expressions of uncertainty during free classification in Wave 1 also received more “No Idea” responses in Wave 2 (Spearman’s ρ = 0.21 to 0.34 across tasks, P < 0.005).

### Machine-learned acoustic structure partly converged with human perceptual structure

To test whether human perceptual organization reflected machine-learned acoustic structure, we compared human-derived context-distance matrices with AI-derived distance matrices from ResNet50 embedding centroids. The best valence-optimized model achieved a validation accuracy of 0.92 (mean = 0.89, SD = 0.06), while the best context-optimized model achieved a validation accuracy of 0.38 across the 18 contexts (mean = 0.32, SD = 0.05).

To assess the contribution of highly aversive contexts, we additionally trained models after excluding vocalizations from castration, fighting, restraint, and crushing. For valence scoring, the reduced-training models achieved a maximum accuracy of 0.92 on the full held-out validation set (mean = 0.88, SD = 0.02) and 0.92 when validation was restricted to the remaining contexts only (mean = 0.89, SD = 0.02). For context classification, accuracy on the held-out validation set restricted to the remaining shared contexts reached 0.39 (mean = 0.31, SD = 0.06).

To test whether high presumed-valence classification was specific to the original context-derived valence assignment, we additionally compared the observed model with models trained after randomly permuting positive and negative labels across the context categories while preserving the original positive-to-negative category ratio. In this fixed-split analysis, the observed assignment yielded a balanced validation accuracy of 0.90. Across the 100 null permutations, mean balanced accuracy was 0.79 (range = 0.56–0.94), and five permuted assignments reached or exceeded the observed accuracy (empirical one-sided P = 0.059). The best-performing null assignment achieved a balanced accuracy of 0.94, exceeding the observed model despite assigning several categories contrary to their original presumed-valence labels, including crushing, barren and negative conditioning as positive and positive conditioning as negative. Thus, high binary classification performance was not specific to the original presumed-valence assignment: multiple arbitrary groupings of recording-context categories were similarly or more strongly discriminable from their acoustic structure.

The context-optimized and valence-optimized AI-derived distance matrices were strongly but not perfectly correlated (Mantel r = 0.68, P < 0.0001), indicating overlapping but nonidentical acoustic organization. Both embedding spaces corresponded significantly with human-derived perceptual structure, with consistently stronger correlations for the valence-optimized model (Table 2). Thus, contexts closer together in AI-derived embedding space also tended to be judged as more similar by human participants.

**Table 2.** Correspondence (Mantel correlations) between AI-derived and human-derived perceptual structure. Columns show correlations for embeddings from models optimized for context classification or presumed valence scoring. The context-optimized and valence-optimized distance matrices were also strongly correlated with one another (Mantel r = 0.682). All Mantel tests had P < 0.0001.

| Comparison | Context model | Valence model |
| --- | --- | --- |
| Wave 1 free classification | 0.598 | 0.735 |
| Wave 2 context classification | 0.518 | 0.724 |
| Wave 2 valence scoring | 0.585 | 0.742 |

When mapping the AI-derived acoustic structure and comparing this to the valence scores assigned to the same sounds by human raters, vocalizations from high-intensity aversive contexts occupied regions associated with more negative human valence ratings (Fig. 5). However, this negative rating structure was absent for several other contexts with presumed negative valence. This was confirmed in a reduced analysis excluding high-intensity aversive vocalizations, in which human-rated valence showed little remaining structure across the AI-derived embedding space (Fig. S12).

**Figure 5.**
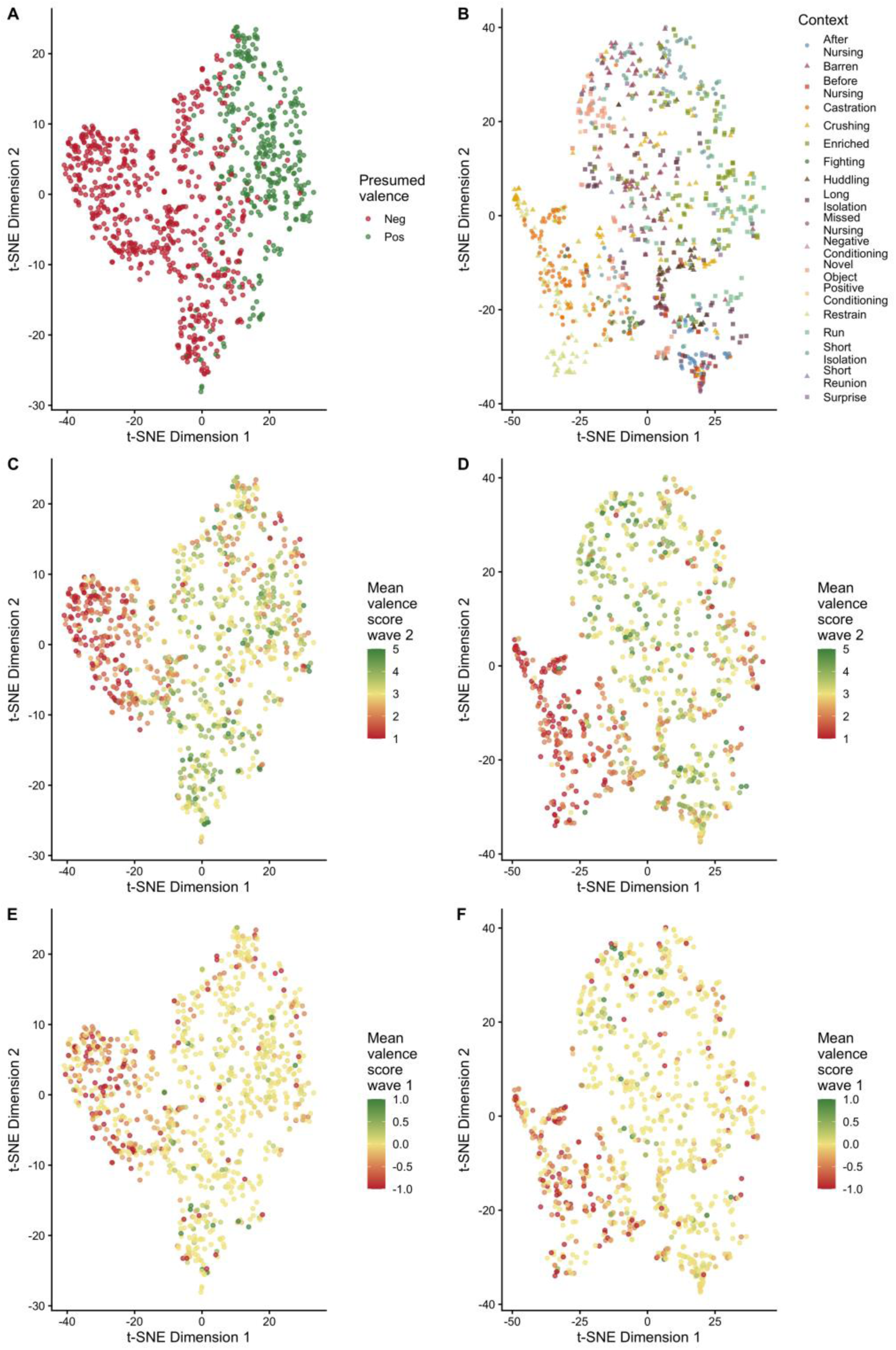
Low-dimensional projections of convolutional neural network embeddings for pig vocalizations. t-distributed stochastic neighbor embedding (t-SNE) projections of ResNet50 embedding vectors from models fine-tuned for valence scoring (A, C, E) or context classification (B, D, F). Each point represents one of the 968 vocalizations scored in both Wave 1 and Wave 2. Points are colored by presumed binary valence in A, recording context in B, mean explicit Wave 2 valence rating in C and D, and mean inferred Wave 1 valence rating in E and F.

### Forced-choice context classification was above-chance but weak

The exact 18-context classification accuracy in Wave 2 was 8.0% when “no idea” responses were retained and 9.5% after excluding “no idea” responses. This exceeded the empirical chance baseline of 5.6% in a participant-wise permutation test, but agreement between source context and selected context was slight (Cohen’s κ = 0.042). Recall was highest for restraint (22.6%) and castration (16.4%), but even these contexts were identified only weakly (Fig. 6A, Table S4). The full confusion matrix showed that classification errors were structured rather than uniformly distributed (Fig. S13). Thus, forced-choice labels detected some structure but did not reliably recover the original recording contexts. A supporting binomial mixed-model analysis found little evidence that participant-related predictors, including pig expertise, explained context-classification accuracy (see Supplementary Text).

**Figure 6.**
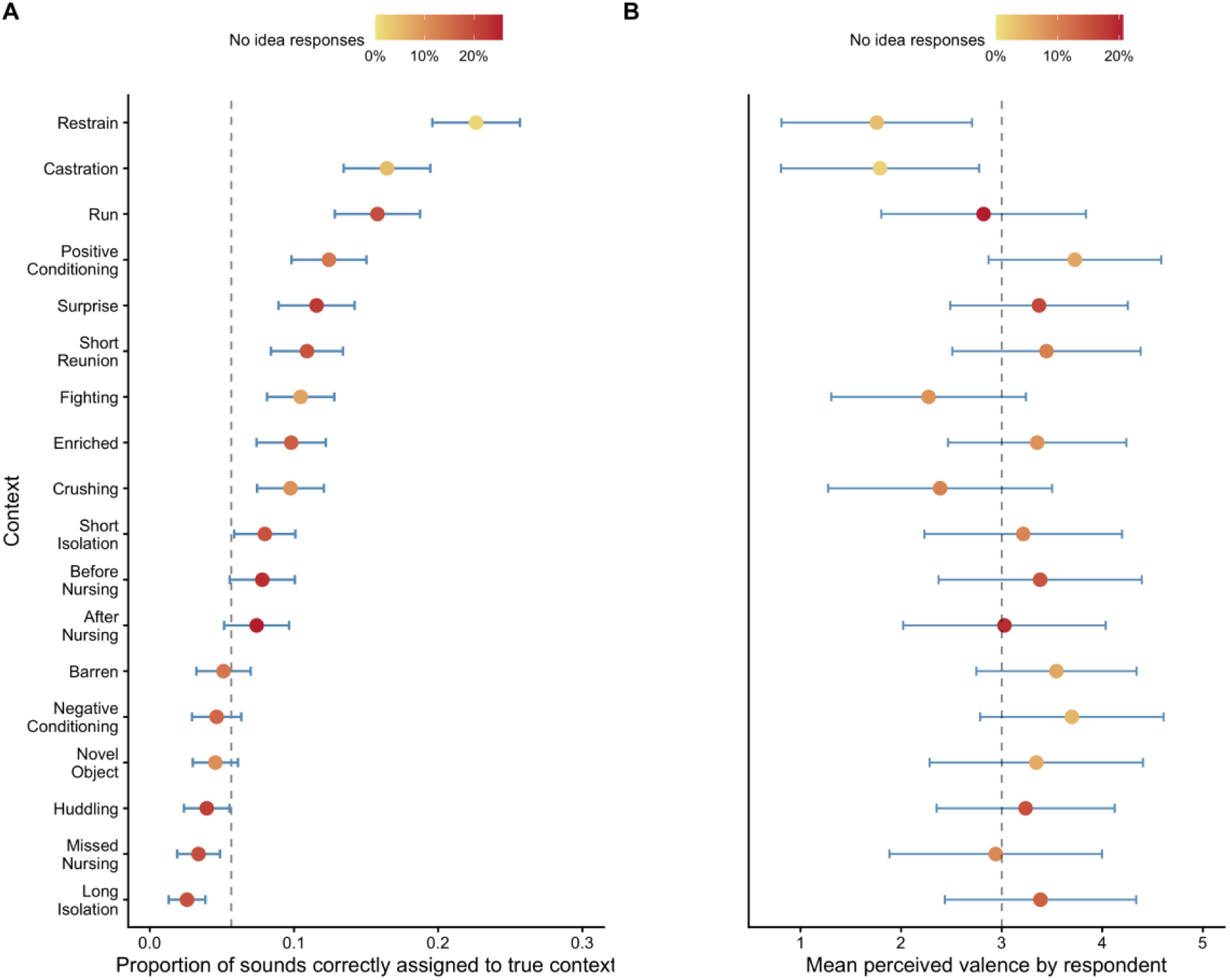
Context-level performance in forced-choice classification. (A) Recall for each recording context in the forced-choice context-classification task. Points show mean recall per context and horizontal bars indicate standard errors. Points are also colored by percentage of “No idea” responses. The vertical dashed line indicates the permutation-based chance baseline. Detailed values are given in Table S3. (B) Mean perceived valence assigned to vocalizations from each recording context on a five-point scale, from very negative (1) to very positive (5). Points show context means and horizontal bars indicate uncertainty around the mean. The vertical dashed line marks the neutral midpoint of the scale. Contexts are ordered based on proportion of correctly classified to the true context as defined by Briefer et al. (2022a).

Duplicate trials showed moderate within-participant consistency: identical context labels were assigned across repeats in 42.5% of trials, with higher consistency for the duplicated castration sound than for the duplicated barren sound (52.8% versus 32.1%). Duplicate-trial recall was also higher for castration than barren context (56.3% versus 7.2%).

### Forced-choice valence scoring accuracy was higher than context classification but driven by high-intensity aversive contexts

Valence ratings showed clearer structure than context classifications but did not cleanly reproduce presumed valence. “No Idea” responses accounted for 9.2% of valence scores and, together with duplicate trials, were excluded from primary analyses. Directional valence accuracy was 42.5% when neutral responses were treated as nonmatching and 60.1% after excluding neutral responses, exceeding the participant-wise permutation baseline of 49.9%. Precision was highest for “Very negative” and “Negative” responses (88.1% and 69.6%; Table S4) and below chance-levels for “Very positive” and “Positive” responses (46.9% and 47.0%; Table S4). Weighted ordinal agreement was low but significant (κ = 0.18), and ratings discriminated presumed positive from presumed negative sounds only modestly (AUC = 0.609 including neutral responses; AUC = 0.634 excluding neutral responses).

Duplicate trials showed higher internal consistency for valence than context labels, with exact and directional valence labels repeated in 77.9% and 84.3% of duplicate trials, respectively. A supporting binomial mixed-model analysis similarly indicated that directional valence accuracy was strongly context-dependent, but showed little evidence that participant-related predictors, including pig expertise, explained accuracy (see Supplementary Text).

The valence confusion matrix showed substantial overlap between presumed positive and presumed negative sounds, with frequent neutral responses (Fig. S14). Perceived valence varied strongly by recording context (Fig. 6B): restraint, castration, fighting, and crushing were rated most negatively, whereas several other contexts presumed as negative were rated near neutral or positive. After excluding high-intensity aversive contexts, weighted ordinal agreement between presumed and perceived valence was no longer different from zero (κ = −0.02, P = 0.24; Fig. S15). Thus, forced-choice valence rating was more informative than exact context classification, but its alignment with presumed valence was driven largely by high-intensity aversive contexts.

### Acoustic correlates of human and neural-network similarity structure

The four conventional acoustic measures showed substantial differentiation among recording contexts (Fig. S16). Consistent with this, the combined acoustic distance structure was positively associated with all human and neural-network similarity structures (Mantel r = 0.65–0.81, all permutation P < 0.001; Table S5). Among individual acoustic measures, median spectral frequency showed particularly strong correspondence with the human similarity structure.

Acoustic characteristics were also associated with explicit human valence judgments from Wave 2. Across sequential cumulative-link mixed models, longer duration and higher median spectral frequency were consistently associated with more negative valence ratings, including after adjustment for listener, animal, recording, and stimulus characteristics (Fig. S17; Table S6). In the most extensively adjusted model, the odds ratios per standard-deviation increase were 0.59 for duration (95% CI, 0.50–0.70) and 0.52 for median spectral frequency (95% CI, 0.43–0.63; both FDR-adjusted P < 0.001). Associations with amplitude-modulation rate did not persist after adjustment for animal characteristics, and Wiener entropy showed no association with perceived valence.

## Discussion

This study showed that pig vocalizations contain reproducible acoustic structure that is recovered similarly across human classification tasks and neural-network representations. Human judgments were also highly consistent across free-classification and explicit-rating tasks, showing that listeners responded systematically to this structure. However, the affective meaning they assigned to vocalizations corresponded only partly with the presumed-valence labels of the source dataset (Briefer et al., 2022a). Agreement was strongest for vocalizations from highly aversive situations and was no longer detectable when these contexts were excluded. Neural networks, by contrast, continued to classify presumed valence with high accuracy, but the original valence grouping did not significantly outperform alternative arbitrary valence groupings of the same recording contexts. Together, these findings show that pig vocalizations contain robust and recoverable acoustic information, while leaving greater uncertainty about what that information represents biologically. High classification accuracy can establish that a grouping is acoustically discriminable, but not that the biological meaning assigned to that grouping has been validated.

### Acoustic and perceptual structure was reproducible across tasks

The first clear pattern was that pig vocalizations contained reproducible structure across different task formats. Similarity patterns derived from free classification, forced-choice context classification, and valence ratings were strongly correlated, showing that listeners organized the vocalizations similarly even when the task changed. This extends previous studies using predefined context or valence categories (Tallet et al., 2010; Greenall et al., 2022) by showing that related perceptual organization also emerges when listeners are given no predefined categories. Human similarity structure also corresponded closely with neural network embeddings and with conventional acoustic structure (Table 2, Fig. S16). Duration and median spectral frequency showed particularly consistent associations, although these analyses do not establish that humans or models relied directly on these features.

The recovered structure was nevertheless broad and strongly overlapping rather than organized into discrete recording contexts or presumed valence categories. The clearest clustering occurred among vocalizations from high-intensity aversive situations (castration, restraint, fighting and crushing) which also elicited the greatest perceptual agreement among listeners. This prominence of highly aversive contexts is consistent with earlier human classification studies of pig vocalizations (Tallet et al., 2010). Outside these contexts, however, similarities were more graded, and vocalizations assigned different presumed valences frequently overlapped in perceptual and acoustic space (Fig. 5). Thus, our findings reinforce that pig vocalizations contain acoustically discriminable information (Briefer et al., 2022a), while showing that this information is not organized cleanly according to recording context or presumed valence.

### Human affective judgments were systematic and reproducible

The free-classification task showed that affective meaning was not the dominant basis on which listeners spontaneously organized pig vocalizations, and only 26.3% of labels were feeling- or emotion-related. This does not imply that the vocalizations lacked affective information or that listeners failed to perceive it. Rather, it shows that affective interpretation was not the dominant response when listeners were free to organize the sounds. When affective terms were used, they were predominantly negative. In Wave 2, however, explicitly asking listeners to rate valence produced more structured judgments, consistent with previous studies using predefined context categories or valence ratings (Tallet et al., 2010; Greenall et al., 2022). The contrast between the two tasks therefore shows that listeners can make systematic valence judgments, even though valence is not the dominant way they spontaneously organize these vocalizations.

Importantly, the valence judgments given across tasks were not arbitrary. Vocalizations that received more positive spontaneous labels in Wave 1 tended to receive more positive ratings in Wave 2 (Fig. 4). This relationship strengthened for vocalizations evaluated by more listeners and remained substantial after excluding the four high-intensity aversive contexts (Fig. S11). Thus, although affective categories were used relatively infrequently in free classification, the emerging affective evaluations were reproducible across participant samples and task formats. This suggests that valence constitutes a systematic dimension of human interpretation of individual vocalizations, even though it is not the primary dimension along which listeners spontaneously organize the vocalization space. Uncertainty showed a similar cross-task consistency, with sounds eliciting more spontaneous uncertainty in Wave 1 also eliciting more ‘No Idea’ responses in Wave 2 indicating that some vocalizations were consistently more difficult to interpret.

Professional background further clarified this distinction between perceptual organization and interpretation. Broad classification structure was highly similar across expertise groups, and professional background did not meaningfully improve forced-choice context classification or valence scoring. It did, however, influence how participants described the sounds. Compared with lay participants, pig farmers and veterinarians used fewer feeling-related labels, animal scientists used more affective terminology, and pig farmers were more likely to use pig-specific descriptors. Tallet et al. (2010) likewise found professional differences in judgments of pig vocalizations, although their pattern differed from ours: pig caretakers rated emotional intensity lower, whereas pig ethologists showed better context recognition than naïve listeners. Such differences have prompted several speculative explanations, including habituation, professional detachment, and greater attention to animal cues, but neither study distinguishes among these mechanisms. Our results therefore provide stronger evidence that professional background affects how pig vocalizations are described than that it consistently changes how accurately their context or affective meaning is recovered.

Different theoretical frameworks offer complementary interpretations of this pattern. The identification–attribution model separates recognition of an observable signal from the later attribution of affective or intentional meaning, whereas the theory of constructed emotion proposes that perception and emotion are shaped together by prior experience and conceptual knowledge (Barrett, 2025; Anikin et al., 2018; Spunt and Lieberman, 2012). The present study cannot distinguish between these accounts or determine the temporal relationship between perceptual organization and affective interpretation. Our study does show that listeners can converge in how pig vocalizations are organized while assigning different meanings to them, and that explicitly asking for affective judgements changes how this shared structure is expressed.

### Human judgments showed limited correspondence with presumed valence

The consistency of human valence judgments contrasted with their much weaker correspondence with the presumed valence assigned to the recording contexts by Briefer et al. (2022a). This correspondence was strongly asymmetric, with negative judgments aligning more often than positive judgments. The clearest agreement was concentrated in vocalizations from castration, restraint, fighting, and crushing. Even within these highly aversive contexts, however, individual vocalizations were not uniformly perceived as negative. Some vocalizations from crushing, for example, received positive mean valence ratings (Fig. 5). Strong agreement at the context-level therefore did not imply that every vocalization produced within that context carried the same perceived valence. More importantly, once these four high-intensity aversive contexts were excluded, correspondence with the presumed-valence labels was no longer detectable in either spontaneous valence-related labels or explicit valence ratings. The apparent human ability to recover the presumed positive-negative distinction was therefore largely driven by a small set of highly aversive situations rather than expressed consistently across the broader vocalization set.

Several non-exclusive explanations could account for this mismatch. Human listeners may respond to affectively relevant variation that is represented imperfectly when a single valence is assigned to an entire recording context, as suggested by positively rated vocalizations from the crushing context. Alternatively, listeners may apply general acoustic rules when judging affect, producing consistent evaluations that do not necessarily reflect the pig’s affective state. Humans have been shown to use similar acoustic cues when judging affective content in conspecific and heterospecific vocalizations (Faragó et al., 2014), which may support cross-species judgments but can also introduce systematic bias. Kelly et al. (2017), for example, found that listeners relied strongly on pitch when judging distress in human and non-human primate infant calls, causing the particularly high-pitched calls of bonobos to be rated as highly distressed across stress-intensity contexts. Consistent human judgments should therefore be taken as evidence that particular acoustic properties reliably shape human affective evaluations, but not by themselves as evidence that these evaluations accurately represent the affective state of the vocalizing animal.

There is currently no independently validated animal-level criterion establishing the momentary affective valence associated with individual pig vocalizations. Context-derived valence labels are themselves inferential, and the acoustic features associated with those labels vary across call types, contexts, and biological conditions. Call duration shows the most recurrent pattern, with longer calls reported in several contexts assigned more negative valence (Tallet et al., 2013; Briefer et al., 2019; Friel et al., 2019; Villain et al., 2020a). Yet, even this relationship varies with call type, anticipated reward, development, and physiological state (Villain et al., 2020b; Nääs et al., 2026), and duration did not consistently covary with peripheral endocrine stress responses in Schräder and Todt (1998). Other acoustic dimensions show even greater heterogeneity, including null, call-type-dependent, and opposing associations with presumed valence (Linhart et al., 2015; Maigrot et al., 2018; Briefer et al., 2019; Friel et al., 2019; Briefer et al., 2022a). The existing literature therefore does not provide a validated acoustic signature against which the affective valence of an individual vocalization can be judged as correct or incorrect.

### Machine-learning discrimination was not specific to presumed-valence labels

The machine-learning analyses sharpened the contrast between acoustic discriminability and human affective interpretation. The valence-optimized neural network discriminated the presumed-valence labels with high within-dataset accuracy (0.92), and performance remained essentially unchanged when castration, restraint, fighting, and crushing were excluded from model training. Human judgments showed the opposite pattern. After excluding the same highly aversive contexts, agreement between perceived and presumed valence was no longer different from zero, and human-rated valence showed little remaining structure across the neural-network embedding space. Thus, the presumed-valence labels remained acoustically discriminable to the neural network even when their correspondence with human-perceived valence had largely disappeared.

The context-category permutation analysis further clarified what this high classification performance establishes. Random guessing would yield a balanced accuracy of 0.50, but this is a weak null when the underlying recording contexts are already acoustically distinct. We therefore randomly reassigned whole context categories to positive and negative labels while preserving the original category ratio, retaining the acoustic structure within each context. Across these arbitrary groupings, mean balanced accuracy reached 0.79. The original presumed-valence assignment achieved 0.90 and was more discriminable than most alternatives, but five of 100 null assignments reached or exceeded this value (empirical one-sided P = 0.059). Hence, classification using the presumed-valence assignment of Briefer et al. (2022a) did not significantly outperform this structured permutation null at α = 0.05.

Thus, high within-dataset classification accuracy was not specific to the affective interpretation attached to the labels. Instead, the recording contexts contained enough systematic acoustic structure for the neural network to discriminate several alternative binary groupings of them. This reflects a broader problem in machine learning, where high predictive performance can arise from features associated with the target labels without establishing that the model has recovered the construct those labels are intended to represent (Geirhos et al., 2020). Evidence of generalization beyond the training animals provides a stronger test. Nääs et al. (2026) achieved 60.9% balanced accuracy across distress and baseline conditions in previously unseen pigs, although performance varied across conditions and growth phases. Such results strengthen evidence that condition-related acoustic information can generalize across animals, but they do not by themselves establish that affective valence is the biological dimension underlying that information.

### Biological Interpretation

This distinction raises a broader biological question: if recoverable acoustic structure cannot be uniquely attributed to valence, what might that structure represent? Evolutionary accounts of animal communication emphasize that vocal signals are shaped by their effects on receivers and need not directly reflect a single internal state (Seyfarth and Cheney, 2003). Accordingly, similar acoustic patterns may arise from different underlying processes, while similar internal states may be expressed differently across contexts, consistent with principles of degeneracy and context dependence (Barrett, 2025; Gendron et al., 2014). Vocalizations may therefore contain affective information without affective valence being the only, or necessarily dominant, dimension organizing their acoustic structure.

Feeding-related vocalizations illustrate this context dependence particularly well. Food anticipation has been treated as a positive context in several pig-vocalization studies (Imfeld-Mueller et al., 2011; Reimert et al., 2013; Leliveld et al., 2016; Maigrot et al., 2018; Briefer et al., 2019, 2022a; Friel et al., 2019), whereas Nääs et al. (2026) operationalized hunger as a negative condition. Although these states are not equivalent, the contrast shows how affective interpretation depends on the specific motivational and experimental context. The same is true for particular types of vocalizations. Vandermeulen et al. (2015), for example, noted that screams may occur during feeding competition without indicating a serious stressful situation, whereas screams occurring at night may carry a very different implication. A vocalization therefore cannot be assigned a fixed affective or welfare meaning independently of the context in which it occurs. The reverse pattern is also possible. Villain et al. (2020b) found different acoustic trajectories during anticipation of two situations both treated as positive, showing that a shared presumed valence need not produce a common acoustic pattern.

Several biological processes may contribute to this non-unique mapping. Nääs et al. (2026) showed substantial developmental variation in pig vocal acoustics, including declining call duration with age, demonstrating that features used to discriminate experimental conditions can also vary independently of valence. Illmann et al. (2013) similarly found that younger and lighter piglets produced more intense vocalizations during simulated crushing, whereas these indicators of vulnerability did not predict vocal intensity during isolation. Other influences, including arousal, motivation, communicative function, and physiological constraints on vocal production, remain plausible contributors whose relative importance is not yet resolved (Villain & Briefer, 2026). Nääs et al. (2026) proposed that context-related changes in respiration and phonation may contribute to acoustic differences under thermal stress, although these pathways were not directly tested. Acoustic structure may therefore reflect several interacting biological dimensions, so neither a particular acoustic feature nor its successful classification is sufficient to identify affective valence.

### Welfare implications

In this study, vocalizations from crushing, restraint, fighting, and castration were among the most consistently distinguished sounds. Vocal monitoring may therefore be particularly useful for detecting specific, acute conditions that produce pronounced acoustic changes. Such detection is welfare-relevant when the adverse consequences of the condition are established independently of the vocal signal, such as mortality, morbidity, tissue damage, injury, physiological disturbance, or impaired access to required resources (Restrepo et al., 2026; Reza et al., 2025). Here, vocalizations serve as indicators of an independently established welfare problem rather than as evidence from which welfare is inferred. This differs fundamentally from using acoustic discrimination to infer a latent affective state and then using that inferred state to support the welfare claim.

For applied welfare monitoring, a stronger validation target is whether detection enables an effective intervention that benefits the animal. Vandermeulen et al. (2015) anticipated this logic by arguing that screams should be interpreted in relation to production context. Detection is therefore most useful when the condition is actionable, and intervention improves outcomes measured at the level of the animal. Such outcomes provide a more direct and testable link between vocal detection and welfare than inferring welfare from an acoustic pattern to a presumed affective state.

This does not imply that subjective experience is irrelevant to welfare or that affective interpretations of vocalizations are necessarily incorrect. Rather, the welfare value of a monitoring system does not need to depend on an additional inference about latent affective state when its benefit to the animal can be demonstrated independently. Following Dawkins (2008), this approach grounds welfare assessment in animal health and in evidence derived from animals’ own choices and behavior. Such evidence is particularly important when welfare assessment aims to place the animal’s perspective at its center (Rault et al., 2025). Predefining that perspective through human-derived affective ratings risks replacing animal-derived evidence with a researcher-imposed interpretation of what the animal experiences. For applied monitoring purposes, vocal signals can thus be valuable without first being validated as readouts of a specific affective state, provided that the conditions they detect and the benefits of responding to them are independently established.

### Limitations

Several limitations should be considered. The classification tasks were challenging, as reflected by substantial participant attrition (50% in Wave 1 and 43% in Wave 2; see Supplementary text) and reports of task difficulty. The final sample may therefore have been enriched for individuals who were particularly motivated to engage with ambiguous vocalizations or interested in questions concerning animal communication and perhaps affect. In Wave 1, the inclusion of a dolphin vocalization as an attention check may also have made non-pig or other-animal interpretations more salient, potentially contributing to participant-defined labels referring to other animals or to sounds judged as “not pig”. Because participation was anonymous, we could not determine the extent of overlap between Wave 1 and Wave 2 samples. Recruitment was also concentrated in Western European countries, likely resulting in a predominantly Western, educated, industrialized, rich, and democratic (WEIRD) sample and limiting generalizability to other populations. This is particularly relevant from a constructionist perspective, because culturally learned concepts may shape how emotion is categorized and attributed across populations.

Further, the vocalizations were derived from a pre-existing dataset in which vocalizations were manually segmented into individual vocal units (Briefer et al., 2022a). In natural settings, however, pigs may produce sequences of acoustically varying calls, such that information is conveyed not only by individual vocalizations but also by their ordering and composition within a vocal bout. Presenting isolated vocal units may therefore have removed sequential information relevant to their perception and interpretation. Because sounds also needed to be audible to human participants, we applied duration and loudness filters that, together with the composition of the source dataset, resulted in unequal numbers of eligible vocalizations across recording contexts (Table S7). Acoustic characteristics were further confounded with recording context, recording conditions, and age, limiting inference about the biological sources of the observed variation. For example, because age and context were not independently varied, age-related acoustic differences may have contributed to some context-level similarities. The context and presumed-valence categories were likewise inherited from the source dataset and reflect prior theoretical interpretations that may not correspond directly to the perceptual organization of the vocalizations. Finally, our design intentionally removed behavioral and environmental context to isolate information available from the acoustic signal itself. The findings therefore concern what can be recovered from isolated vocalizations, rather than the full set of cues available during natural communication.

## Conclusion

The findings from this study have important implications for the use of animal vocalizations in welfare research. Vocal features may vary systematically across affective conditions under controlled experimental designs, but our results show that these differences do not translate consistently into discrete, context-independent affective meaning. Instead, human listeners recovered reproducible acoustic structure, yet this structure overlapped across contexts and corresponded unevenly with presumed valence. Neural networks likewise achieved high classification accuracy, but comparable performance for several arbitrary context groupings showed that this accuracy was not specific to the presumed-valence distinction. Pig vocalizations therefore contain rich and recoverable information, but acoustic discriminability alone cannot establish its affective or welfare meaning. Future work should distinguish successful detection from biological interpretation and independently establish what the detected vocal variation represents for the animal.

## Materials and Methods

### Study design

We conducted a two-wave online classification study to examine how human listeners organize isolated pig vocalizations in the absence or presence of predefined categories. Wave 1 used a free-classification task, in which participants grouped vocalizations according to perceived similarity and provided their own category labels. Wave 2 used forced-choice classification tasks, in which a separate participant sample classified vocalizations into predefined recording-context categories and subsequently rated the same vocalizations on an explicit valence scale. These human-derived response structures were compared with acoustic structure learned by convolutional neural network models trained on spectrograms. The analyses were designed to test whether isolated pig vocalizations support context-specific classification or valence-specific scoring, and whether human perceptual structure converges with machine-learned acoustic structure.

The study was preregistered before analysis at https://doi.org/10.17605/OSF.IO/EJUR7. The study followed the preregistered overall two-wave design comparing free and predefined-category classification of context-free pig vocalizations. Several procedural details differed from preregistration: Wave 2 used 25 rather than 20 sounds, used an 18-context classification task followed by a five-point Likert scale valence task, and did not include a non-pig outlier. Analyses of human classification patterns reflect the preregistered aims, whereas the semantic-label analyses, neural-network embedding analyses, human–machine correspondence tests, and sensitivity analyses are reported as exploratory or sensitivity analyses.

### Source vocalization dataset and stimulus selection

Pig vocalizations were obtained from the Soundwel database (Briefer et al., 2022b), containing recordings collected from eighteen experimentally defined contexts in domestic pigs, each previously assigned a presumed context-derived valence (negative or positive) including both low- and high-frequency calls (Briefer et al., 2022a). From the 6,887 recordings, a subset of 3,140 vocalizations was selected based on objective acoustic quality criteria. Recordings were filtered using automated signal processing implemented in a custom Python script. Files were excluded if they showed low signal-to-noise ratio (<6 dB), short duration (<0.10 s), or insufficient loudness (LUFS < −30), ensuring adequate perceptual quality for human listeners. An overview of the number of sounds per context after filtering is provided in Table S7.

### Structured inferential audit of prior pig-vocalization evidence

We conducted a structured inferential audit of previous studies linking pig vocalizations to affective state, valence, arousal, stress, or welfare. The aim was to distinguish evidence that vocalizations discriminate experimental or observational conditions from evidence that such differences specifically reflect affective state. Studies were evaluated against five auxiliary assumptions concerning construct realization (A1), construct comparability (A2), construct specificity (A3), measurement correspondence (A4), and transportability (A5). For each study, the senior author (C.W.) assessed the relevant operationalization, supporting evidence, and inferential limitations using the full article and supplementary materials where available.

Each assumption was classified as Direct, External, Inconclusive, Contrasted, or Not tested, and the highest supported claim was assigned from C0 (condition discrimination) to C4 (prediction of affective valence in a novel context). Coding was conservative: studies were not assigned a higher claim level when this required unresolved auxiliary assumptions. Reuse of datasets, inherited affective labels, and reliance on previously proposed acoustic-affect mappings were also recorded as sources of evidential dependence. Summary classifications are presented in Table 1; the complete study-level evidence and rationale for each classification are provided in Table S1.

### Participants and data collection procedures

The study was approved by the ETH Zurich Ethics Committee (number 26 ETHICS-004). Data was collected online in two waves, and participants provided informed consent prior to participation. Screenshots of both surveys are provided in Fig. S18. The surveys were provided in English, Dutch, French, German, Italian and Spanish. In total, 448 participants started the survey in Wave 1 and 224 completed it; in Wave 2, 374 participants started the survey and 159 completed it. Incomplete registrations were retained in the dataset but excluded from all analyses.

In Wave 1 (February 2026), participants classified pig vocalizations in a free classification task with 40 vocalizations: 2 vocalizations randomly drawn from each of the 18 context categories (18x2=36), 2 outliers (duplicated dolphin sound) and 2 duplicates from the castration context. Duplicate and outlier sounds were included as internal controls to assess participant attention and categorization consistency. Participants freely grouped vocalizations into self-defined categories based on perceived similarity. No constraints were imposed on the number or naming of categories.

In Wave 2 (April 2026), participants classified pig vocalizations in a forced-choice classification and scoring task with 25 vocalizations: 1 vocalization randomly drawn from each of the 18 context categories, 2 duplicates from the castration and the barren context (2x2=4) and 3 randomly drawn vocalizations. Wave two consisted of two parts. In a first part, participants needed to classify the 25 sounds in the 18 predefined context categories or in a “no idea” category. The order of context categories was randomized per participant. Next, respondents scored the same 25 sounds according to their perceived positivity or negativity on a 5-point Likert scale (“very negative”, “negative”, “neutral”, “positive” or “very positive”) or in a “no idea” category. Respondents were not informed in advance that the same 25 sounds were used in both tasks of Wave 2.

For both waves, vocalizations were randomly drawn per context from the selected subset of 3,140 vocalizations. Vocalizations were shown simultaneously in a randomized order and participants needed to drag these to either self-created categories (Wave 1) or in forced-choice categories (Wave 2). The number of times participants played each sound was recorded. After the classification task, participants were asked to provide basic personal information (profession, gender, age group) and self-assessed expertise with pigs on a 0 to 10 scale. As identifying information was not registered, it is unknown how many respondents filled surveys from both waves.

### Statistical analysis overview

All analyses were conducted in R version 4.5.2 unless otherwise specified. Raw JSON data from the online survey platform were parsed and converted into sound-level response tables. Analyses were conducted at three main levels: individual sound presentations, participant-defined categories, and context-level similarity matrices. The main human-response outcomes were free-classification co-classification similarity in Wave 1, semantic content of participant-defined labels in Wave 1, forced-choice context classification in Wave 2, forced-choice valence ratings in Wave 2, and correspondence between human-derived and machine-learned context-distance matrices.

### Free-classification similarity and clustering

For Wave 1, within-context consistency was quantified as the proportion of participants who placed the two vocalizations from the same recording context into the same self-defined category. To quantify perceived similarity between contexts, participant groupings were converted into context-level co-classification matrices. For each participant, context pairs were scored according to whether vocalizations from the two contexts were placed in the same participant-defined category. Participant-level matrices were then averaged to obtain a group-level similarity matrix, with each cell representing the proportion of participants who grouped sounds from the two contexts together. Dissimilarity matrices were calculated as 1 − similarity.

The group-level dissimilarity matrix was used for clustering and visualization. Principal coordinates analysis (PCoA), equivalent to classical multidimensional scaling, was applied to the dissimilarity matrix to obtain a two-dimensional representation of the perceptual structure among recording contexts. The same approach was also used for expertise-stratified matrices to test whether broad perceptual structure differed between participant groups.

### Semantic analysis of free-classification labels

Participant-defined category labels from Wave 1 were cleaned and standardized before analysis. Capitalization, punctuation, and emoji characters were removed for text-based analyses while preserving semantic content. Non-English responses were translated to English using Google Translate API accessed through R. Translated labels were tokenized, lemmatized, and part-of-speech tagged using the udpipe English model. Lemmas were assigned to semantic categories using WordNet through Python’s NLTK interface in R.

The semantic analysis focused on five non-exclusive categories: feeling, state, event, animal, and emotion. These categories were derived from WordNet semantic domains rather than assigned manually. In descriptive terms, “feeling” referred to internal sensations or subjective perceptions; “state” referred to conditions or states of being; “event” referred to actions or occurrences; “animal” referred to words denoting animals or animal-related terms; and “emotion” referred to explicit emotion concepts. Each lemmatized term was mapped to one or more WordNet lexical domains, with domains relevant to the present study prioritized when multiple classifications were available. For adjectives, the semantic domain of a derivationally related noun was used where possible; for example, *angry* was linked to *anger* and classified within the feeling domain. Thus, “feeling” and “emotion” represented distinct WordNet-derived semantic categories rather than an a priori distinction imposed by the researchers. Labels could be assigned to multiple categories, allowing overlap between semantic types. Analyses were conducted at both the label level and the participant level.

Feeling- and emotion-related labels were additionally coded for inferred valence. Labels with clearly positive meaning were manually coded as +1, labels with clearly negative meaning were coded as −1, and labels that were unclear, mixed, neutral, or not feeling- or emotion-related were coded as 0. For each sound, these label-level values were averaged across participants to obtain a Wave 1 inferred-valence proxy. This proxy was used only as an index of human interpretation of the sound and was not treated as a direct measure of the animal’s affective state.

Agreement between the Wave 1 inferred-valence proxy and Wave 2 explicit valence ratings was assessed using Spearman rank correlations across sounds rated in both waves. Analyses were repeated using increasingly strict inclusion thresholds based on the minimum number of raters per sound in both waves. These thresholds were used to evaluate whether the association between Wave 1 and Wave 2 valence estimates depended on sparse sound-level sampling.

As a parallel analysis, participant-generated labels were also coded for spontaneous expressions of uncertainty. The primary, strict definition included labels explicitly indicating inability or uncertainty in classifying the sound (e.g., “no idea,” “don’t know,” “unknown,” or “too short to judge”). A broader sensitivity definition additionally included labels containing a question mark, which had been converted during text cleaning to an “uncertainty” token. For each sound, the proportion of Wave 1 participants expressing uncertainty was calculated. These sound-level proportions were compared with the proportions of Wave 2 participants selecting “No Idea” in the forced-choice context and valence tasks using Spearman rank correlations. Analyses were repeated using minimum inclusion thresholds of one to four raters per sound in both waves. P values for the four uncertainty comparisons within each rater threshold were adjusted using the Holm procedure.

### Forced-choice context classification

For the Wave 2 context-classification task, accuracy was calculated at the individual sound-response level. A response was coded as correct when the participant-selected context matched the recording context of the sound. Duplicate presentations were removed to calculate classification metrics, as they might inflate results. Overall accuracy was calculated as the proportion of correct responses. Precision was calculated as the proportion of responses assigned to a given context that corresponded to sounds truly originating from that context. Context-specific recall was calculated as the proportion of sounds from a given recording context that were correctly assigned to that context. F1 scores were calculated as the harmonic mean of recall and precision. The proportion of “no idea” responses was calculated separately for each true recording context.

Context-classification performance was evaluated against a chance baseline. A participant-wise permutation test was performed by shuffling context responses within participant and recalculating overall accuracy across 1,000 permutations. Agreement between true and assigned context labels was also quantified using Cohen’s kappa after excluding “no idea” responses.

### Forced-choice valence scoring

For Wave 2 valence scoring, ordinal valence responses were coded from −2 to +2. “No idea” responses were excluded from analyses requiring numerical valence scores and summarized separately as an uncertainty measure. Duplicate presentations were removed to calculate classification metrics, as they might inflate results. Mean explicit valence was calculated for each sound by averaging numerical valence responses across participants. Context-level valence summaries were calculated by averaging sound-level means within recording context.

To evaluate whether explicit valence ratings aligned with the presumed binary valence labels from the source dataset, responses were summarized as negative, neutral, or positive where appropriate. “Very negative” and “negative” responses were treated as negative, “positive” and “very positive” responses were treated as positive, and “neutral” responses were retained as neutral. Classification metrics were calculated for presumed negative and presumed positive source labels, with “no idea” responses excluded from accuracy calculations but reported separately.

### Forced-choice similarity matrices

To compare Wave 2 forced-choice tasks with Wave 1 free classification, context-level similarity matrices were also constructed from Wave 2 responses. For the context-classification task, two recording contexts were considered similar when participants assigned sounds from those contexts to the same predefined context category. For the valence-scoring task, two recording contexts were considered similar when participants assigned sounds from those contexts to the same valence category. Participant-level matrices were averaged to obtain group-level similarity matrices, and dissimilarities were calculated as 1 − similarity. These matrices were compared with the Wave 1 free-classification matrix using Mantel tests.

### Duplicate and outlier consistency

Duplicate and outlier sounds were included as internal controls to assess within-participant response consistency. For each duplicated sound, consistency was defined as whether a participant assigned the two presentations of the same sound to the same category or response option. Mean duplicate consistency was calculated overall and separately for each duplicated sound. Duplicate trials were used for internal consistency analyses and were excluded from context-level performance metrics unless otherwise specified.

### Machine-learning embedding analysis: scope and interpretation

The machine-learning analysis was used to characterize acoustic structure in the vocalization dataset and to compare that structure with human-derived perceptual organization. The validation accuracies of the neural-network models were interpreted as internal discrimination performance within the available spectrogram dataset. They were not treated as evidence that the models recovered animal affective states. The pipeline was based on the ResNet50 spectrogram-classification approach used by Briefer et al. (2022a)

### ResNet50 fine-tuning

Spectrogram images from the source dataset were matched to the metadata table using the spectrogram filename field. The neural-network analyses used vocalizations that were classified by at least one participant in Wave 1 or Wave 2 and had an available spectrogram image. This yielded 2,507 spectrograms for model training and embedding extraction.

All spectrograms were converted to RGB, resized to 224 × 224 pixels, and preprocessed according to the ImageNet ResNet50 convention. Pixel values were kept on the original 0–255 scale, channels were reordered from RGB to BGR, and ImageNet channel means were subtracted. A ResNet50 model pretrained on ImageNet was used as the image backbone with the original ImageNet classification head removed and global average pooling enabled, yielding a 2048-dimensional representation layer. Two task-specific models were trained separately: one optimized for binary presumed-valence classification and one optimized for recording-context classification. As a sensitivity analysis, both models were additionally retrained after excluding vocalizations from the high-intensity aversive contexts of castration, restraint, fighting, and crushing, to assess whether model discrimination and the resulting acoustic structure were driven primarily by these contexts. For each model, the ResNet50 backbone was initially frozen, after which the final residual block, corresponding to layers with names containing conv5, was unfrozen for fine-tuning. A dropout layer with rate 0.5 and a task-specific softmax output layer were added on top of the global-average-pooling output.

For each task, the data were split into training and validation sets using a stratified 70/30 split at the spectrogram level. Training was repeated ten times per task using different random seeds and train-validation splits. Models were trained for 20 epochs with a mini-batch size of 32. Optimization used stochastic gradient descent with momentum 0.9 and categorical cross-entropy loss. The initial learning rate was 0.001 and was multiplied by 10^-0.5^ every 10 epochs. To account for class imbalance, inverse-frequency class weights were applied during training. For each run, the checkpoint with the highest validation accuracy was saved, and the best-performing run for each task was selected for downstream embedding analyses. Because train-validation splitting was performed at the spectrogram level, validation accuracy should be interpreted as within-dataset discrimination performance and not as an estimate of generalization to independent animals or recording conditions.

The reduced models were trained using the same architecture, preprocessing, train-validation splitting procedure, and training parameters as the corresponding full-dataset models. Their validation performance was evaluated both on the full held-out validation set and, where relevant, after restricting validation to the contexts retained in the reduced analysis.

As an additional null-model analysis, we tested whether high presumed-valence classification accuracy was specific to the original context-derived valence assignment. Valence labels were permuted across contexts. The permutation analysis used the 17 source context category levels, which are distinct from the 18 recording-context labels used in the human classification task. The original assignment comprised seven positive and ten negative context categories; each permutation preserved this ratio, and all spectrograms belonging to a given category inherited the same permuted label. We trained 100 models using unique randomly selected alternative label assignments, together with one model using the original assignment. All runs used the same ResNet50 architecture and training parameters and the same fixed 70/30 spectrogram-level train-validation split, generated from the original valence labels, so that only the context-to-valence assignment varied across permutations. Validation balanced accuracy was used as the primary metric because the number of spectrograms assigned to each permuted class varied with category size. The observed balanced accuracy was compared with the permutation distribution using a one-sided empirical P value, calculated as (1 + number of permutation accuracies greater than or equal to the observed accuracy)/(100 + 1).

### Embedding extraction and visualization

After selecting the best valence-optimized and context-optimized models, embeddings were extracted for all retained spectrograms from the global-average-pooling layer. This produced two 2048-dimensional representation spaces: one optimized for presumed-valence discrimination and one optimized for recording-context discrimination. Exact duplicate embedding rows were removed before dimensionality reduction and context-level clustering.

To visualize individual vocalizations, t-distributed stochastic neighbor embedding (t-SNE) was applied separately to the valence-optimized and context-optimized embedding matrices. For the valence-optimized model, t-SNE was run with two dimensions, PCA initialization, perplexity = 50, theta = 0.5, 1,000 iterations, and a fixed random seed. For the context-optimized model, the same settings were used except that perplexity was set to 20. The t-SNE coordinates were calculated on the full retained embedding set; plots used for comparison with human ratings were then restricted to the 968 vocalizations with the required Wave 1 and Wave 2 human-response data.

Embedding-space analyses were also repeated after excluding vocalizations from the four high-intensity aversive contexts, allowing us to assess whether the correspondence between acoustic structure and human-rated valence persisted in their absence.

### Comparison between machine-learned and human-derived structure

For context-level comparisons, embeddings were summarized by recording context. Embedding dimensions with zero variance were removed, and the remaining dimensions were centered and scaled across individual vocalizations. Scaled vocalization-level embeddings were averaged within each recording context to obtain one context centroid per model. Pairwise Euclidean distances between context centroids were calculated to obtain AI-derived context-distance matrices. These matrices were used for principal coordinates visualization, and comparison with human-derived perceptual structure.

AI-derived distance matrices were compared with human-derived distance matrices using Mantel tests. Human similarity matrices from Wave 1 free classification, Wave 2 forced-choice context classification, and Wave 2 forced-choice valence scoring were converted to distance matrices as 1 − similarity before Mantel testing. These analyses tested whether contexts judged as perceptually similar by humans were also close in machine-learned acoustic embedding space.

### Acoustic correlates of human and neural-network similarity structure

To further characterize the acoustic dimensions underlying the recovered similarity structure, we examined four acoustic measures available in the SoundWel dataset and previously selected by Briefer et al. (2022a): call duration, amplitude-modulation rate, median spectral frequency (Q50), and Wiener entropy. Measures were standardized across vocalizations and averaged within recording context, and Euclidean distances were calculated between context-level acoustic centroids. These acoustic distances were compared with the human free-classification, forced-choice context and valence similarity structures, and the context- and valence-trained neural-network representations using Pearson Mantel tests with 9,999 permutations. Associations with individual acoustic measures were examined exploratorily, with false-discovery-rate correction across the four measures.

We additionally examined whether these four acoustic measures were associated with explicit valence ratings in Wave 2 using cumulative-link mixed models. Ratings of “No Idea” were excluded, and acoustic predictors were standardized. Models included random intercepts for participant and vocalization and were built sequentially, starting with the four acoustic measures and subsequently adding listener characteristics (age, gender, pig expertise, profession, and sound-analysis experience), pig characteristics (age category and sex), recording-related variables (loudness, and signal-to-noise ratio), and finally recording context and call type. False-discovery-rate correction was applied across the four acoustic predictors within each model.

### Sensitivity analyses

Sensitivity analyses were conducted to test whether apparent valence structure was driven primarily by high-intensity aversive contexts. Unless otherwise stated, high-intensity aversive contexts were defined as castration, restraint, fighting, and crushing. Key analyses involving inferred valence, explicit valence, and embedding-space visualization were repeated after excluding these contexts. These analyses evaluated whether correspondence between presumed valence, human valence ratings, and acoustic embedding structure persisted when the most salient aversive contexts were removed.

## Acknowledgments

We would like to thank Marleen Berger, Dré Sleurs, Edoardo Henzen, Marie-Emilie Lebachelier de la Rivière, Álvaro López-Valiñas and Linard Balke for testing the online survey and checking the different languages. We gratefully acknowledge Jan Peters, whose participation in the first wave of the study was remembered by his family as a moment of joy and laughter. Moreover, we would like to acknowledge all organizations and people that helped in spreading our surveys.

## Author Contributions

Carmen Winters conceived the study and led the experimental design and setup, with contributions from Wim Gorssen and Ben Sleurs. Ben Sleurs developed and implemented the online survey used for data collection. Wim Gorssen performed most of the statistical and machine-learning analyses. Carmen Winters conducted the semantic language analysis and contributed to the remaining analyses. Carmen Winters and Wim Gorssen wrote the original manuscript draft and revised the manuscript jointly. Ben Sleurs contributed to manuscript revision and reviewed the final version. All authors approved the final manuscript.

## Competing Interest Statement

The authors have no competing interests.

## Funding

Wim Gorssen was funded by a Swiss Postdoctoral Fellowship (Grant Number 234026) of the Swiss National Science Foundation (SNSF). Carmen Winters and Ben Sleurs received no specific funding for this work. The funding body played no role in the design of the study, collection analysis, interpretation of data and in writing the manuscript.

## Data availability statement

The pig vocalizations used in this study were obtained from the Soundwel Database and are publicly available through Zenodo: https://zenodo.org/record/8252482 (Briefer et al., 2022b). The anonymized human participant classifications collected for the present study, including responses for the analyzed subset of vocalizations, are publicly available at https://doi.org/10.17605/OSF.IO/Q89NR (https://osf.io/q89nr/files/osfstorage).

## Code availability statement

All analysis code required to reproduce the results reported in this study are publicly available at https://doi.org/10.17605/OSF.IO/Q89NR (https://osf.io/q89nr/files/osfstorage).

## Notes

https://doi.org/10.17605/OSF.IO/EJUR7

https://doi.org/10.17605/OSF.IO/Q89NR

https://zenodo.org/record/8252482

### Competing Interest Statement

The authors have declared no competing interest.

### Summary of Updates

This version has been substantially revised in response to peer review. The Abstract, Introduction, and Discussion were restructured to clarify the distinction between acoustic discriminability, human affective interpretation, and validation of animal affective state. We added a structured inferential audit of previous studies linking pig vocalizations to affective state and expanded engagement with the relevant literature. The rationale and interpretation of the free classification task were clarified. We also added a context category permutation analysis to test whether high neural network classification performance was specific to the original presumed valence grouping. Additional sensitivity analyses and methodological clarifications were included, and the interpretation of the human and machine learning results was revised accordingly. Figures, tables, references, and Supporting Information were updated to reflect these changes.

## References

1. Anikin, A., Bååth, R., & Persson, T. (2018). Human non-linguistic vocal repertoire: call types and their meaning. Journal of nonverbal behavior, 42(1), 53–80. 10.1007/s10919-017-0267-y

2. Barrett, L. F., Atzil, S., Bliss-Moreau, E., Chanes, L., Gendron, M., Hoemann, K., Katsumi, Y., Kleckner, I. R., Lindquist, K. A., Quigley, K. S., Satpute, A. B., Sennesh, E., Shaffer, C., Theriault, J. E., Tugade, M., & Westlin, C. (2025). The Theory of Constructed Emotion: More Than a Feeling. Perspectives on psychological science : a journal of the Association for Psychological Science, 20(3), 392–420. 10.1177/17456916251319045

3. Bliss-Moreau, E. (2017). Constructing nonhuman animal emotion. Current Opinion in Psychology, 17, 184–188. 10.1016/j.copsyc.2017.07.011

4. Briefer, E. F. (2012). Vocal expression of emotions in mammals: mechanisms of production and evidence. Journal of Zoology, 288(1), 1–20. 10.1111/j.1469-7998.2012.00920.x

5. Briefer, E. F., Vizier, E., Gygax, L., & Hillmann, E. (2019). Expression of emotional valence in pig closed-mouth grunts: Involvement of both source-and filter-related parameters. The Journal of the Acoustical Society of America, 145(5), 2895–2908. 10.1121/1.5100612

6. Briefer, E. F., Sypherd, C. C., Linhart, P., Leliveld, L. M. C., Padilla de la Torre, M., Read, E. R., Guérin, C., Deiss, V., Monestier, C., Rasmussen, J. H., Špinka, M., Düpjan, S., Boissy, A., Janczak, A. M., Hillmann, E., & Tallet, C. (2022a). Classification of pig calls produced from birth to slaughter according to their emotional valence and context of production. Scientific reports, 12(*1*), 3409. 10.1038/s41598-022-07174-8

7. Briefer, E. F., Sypherd, C. C. R., Leliveld, L. M. C., Padilla de la Torre, M., & Tallet, C. (2022b). The Soundwel Database: a labeled pig vocalization repository [Data set]. Scientific Reports, 12(*1*), 3409. 10.1038/s41598-022-07174-8

8. Dawkins, M. S. (2008). The science of animal suffering. Ethology, 114(10), 937–945. 10.1111/j.1439-0310.2008.01557.x

9. Faragó, T., Andics, A., Devecseri, V., Kis, A., Gácsi, M., & Miklósi, Á. (2014). Humans rely on the same rules to assess emotional valence and intensity in conspecific and dog vocalizations. Biology letters, 10(1), 20130926. 10.1098/rsbl.2013.0926

10. Friel, M., Kunc, H. P., Griffin, K., Asher, L., & Collins, L. M. (2019). Positive and negative contexts predict duration of pig vocalisations. Scientific reports, 9(1), 2062. 10.1038/s41598-019-38514-w

11. Geirhos, R., Jacobsen, J. H., Michaelis, C., Zemel, R., Brendel, W., Bethge, M., & Wichmann, F. A. (2020). Shortcut learning in deep neural networks. Nature Machine Intelligence, 2(11), 665–673. 10.1038/s42256-020-00257-z

12. Gendron, M., Lindquist, K. A., Barsalou, L., & Barrett, L. F. (2012). Emotion words shape emotion percepts. *Emotion (Washington*, D.C.), 12(2), 314–325. 10.1037/a0026007

13. Gendron, M., Roberson, D., van der Vyver, J. M., & Barrett, L. F. (2014). Cultural relativity in perceiving emotion from vocalizations. Psychological science, 25(4), 911–920. 10.1177/0956797613517239

14. Greenall, J. S., Cornu, L., Maigrot, A. L., de la Torre, M. P., & Briefer, E. F. (2022). Age, empathy, familiarity, domestication and call features enhance human perception of animal emotion expressions. Royal Society open science, 9(12), 221138. 10.1098/rsos.221138

15. Illmann, G., Hammerschmidt, K., Špinka, M., & Tallet, C. (2013). Calling by domestic piglets during simulated crushing and isolation: a signal of need? . PLoS One, 8(12), e83529. 10.1371/journal.pone.0083529

16. Imfeld-Mueller, S., Van Wezemael, L., Stauffacher, M., Gygax, L., & Hillmann, E. (2011). Do pigs distinguish between situations of different emotional valences during anticipation? . Applied Animal Behaviour Science, 131(3-4), 86–93. 10.1016/j.applanim.2011.02.009

17. Kelly, T., Reby, D., Levréro, F., Keenan, S., Gustafsson, E., Koutseff, A., & Mathevon, N. (2017). Adult human perception of distress in the cries of bonobo, chimpanzee, and human infants. Biological Journal of the Linnean Society, 120(4), 919–930. 10.1093/biolinnean/blw016

18. Lindquist, K. A., Barrett, L. F., Bliss-Moreau, E., & Russell, J. A. (2006). Language and the perception of emotion. Emotion, 6(1), 125–138. 10.1037/1528-3542.6.1.125

19. Linhart, P., Ratcliffe, V. F., Reby, D., & Špinka, M. (2015). Expression of emotional arousal in two different piglet call types. PloS one, 10(8), e0135414. 10.1371/journal.pone.0135414

20. Lefèvre, R. A., Sypherd, C. C., & Briefer, É. F. (2025). Machine learning algorithms can predict emotional valence across ungulate vocalizations. IScience, 28(2), 111834. 10.1016/j.isci.2025.111834

21. Leliveld, L. M., Düpjan, S., Tuchscherer, A., & Puppe, B. (2016). Behavioural and physiological measures indicate subtle variations in the emotional valence of young pigs. Physiology & Behavior, 157, 116–124. 10.1016/j.physbeh.2016.02.002

22. Maigrot, A. L., Hillmann, E., & Briefer, E. F. (2018). Encoding of emotional valence in wild boar (Sus scrofa) calls. Animals, 8(6), 85. 10.3390/ani8060085

23. Mason, G. J., & Mendl, M. T. (2026). Measuring the unmeasurable: The construct validation of affective state indicators. In G. J. Mason, B. L. Nielsen, & M. T. Mendl (Eds.), Assessing animal welfare: A guide to the valid use of indicators of affective states (pp. 26–54). John Wiley & Sons Ltd. 10.1002/9781394182237.ch2

24. Nääs, I. D. A., Pereira, D. F., Cordeiro, A. F. D. S., & Lima, N. D. D. S. (2026). Hierarchical Acoustic Encoding Distress in Pigs: Disentangling Individual, Developmental, and Emotional Effects with Subject-Wise Validation. Animals, 16(8), 1148. 10.3390/ani16081148

25. Nicolaisen, T. J., Bollmann, K. E., Hennig-Pauka, I., & Fischer, S. C. L. (2025). Framework for Classification of Fattening Pig Vocalizations in a Conventional Farm with High Relevance for Practical Application. Animals, 15(17), 2572. 10.3390/ani15172572

26. Rault, J. L., Bateson, M., Boissy, A., Forkman, B., Grinde, B., Gygax, L., Harfeld, J. L., Hintze, S., Keeling, L. J., Kostal, L., Lawrence, A. B., Mendl, M. T., Miele, M., Newberry, R. C., Sandøe, P., Špinka, M., Taylor, A. H., Webb, L. E., Whalin, L., & Jensen, M. B. (2025). A consensus on the definition of positive animal welfare. Biology letters, 21(1), 20240382. 10.1098/rsbl.2024.0382

27. Reimert, I., Bolhuis, J. E., Kemp, B., & Rodenburg, T. B. (2013). Indicators of positive and negative emotions and emotional contagion in pigs. Physiology & behavior, 109, 42–50. 10.1016/j.physbeh.2012.11.002

28. Restrepo, V., Herter, S., & Fischer, S. C. L. (2026). Alarming pig vocalization-based prediction using the self-supervised BEATs model. In J. Dörr et al. (Eds.), Datenräume in der Land-, Forst- und Ernährungswirtschaft: Referate der 46. GIL-Jahrestagung (Lecture Notes in Informatics, pp. 428–434). Gesellschaft für Informatik. 10.18420/giljt2026_54

29. Reza, M. N., Ali, M. R., Haque, M. A., Jin, H., Kyoung, H., Choi, Y. K., Kim, G., & Chung, S. O. (2025). A review of sound-based pig monitoring for enhanced precision production. Journal of animal science and technology, 67(2), 277–302. 10.5187/jast.2024.e113

30. Schrader, L., & Todt, D. (1998). Vocal quality is correlated with levels of stress hormones in domestic pigs. Ethology, 104(10), 859–876. 10.1111/j.1439-0310.1998.tb00036.x

31. Seyfarth, R. M., & Cheney, D. L. (2003). Signalers and receivers in animal communication. Annual review of psychology, 54(1), 145–173. 10.1146/annurev.psych.54.101601.145121

32. Spunt, R. P., & Lieberman, M. D. (2012). An integrative model of the neural systems supporting the comprehension of observed emotional behavior. Neuroimage, 59(3), 3050–3059. 10.1016/j.neuroimage.2011.10.005

33. Tallet, C., Špinka, M., Maruščáková, I., & Šimeček, P. (2010). Human perception of vocalizations of domestic piglets and modulation by experience with domestic pigs (Sus scrofa). Journal of comparative psychology, 124(1), 81–91. 10.1037/a0017354

34. Tallet, C., Linhart, P., Policht, R., Hammerschmidt, K., Šimeček, P., Kratinova, P., & Špinka, M. (2013). Encoding of situations in the vocal repertoire of piglets (Sus scrofa): a comparison of discrete and graded classifications. PloS one, 8(8), e71841. 10.1371/journal.pone.0071841

35. Valfré, L. R., Buchini, J. L. C., Braz, J. M., Malini, Y., Braga, C. S., Da Silva, M. D. A., & Caldara, F. R. (2026). Resilience Metrics in Pigs: How Novelty and Environmental Complexity Shape Affective State and Neurobiological Outcomes. Animals, 16(11), 1742. 10.3390/ani16111742

36. Vandermeulen, J., Bahr, C., Tullo, E., Fontana, I., Ott, S., Kashiha, M., Guarino, M., Moonds, C.P., Tuyttens, F.A., Niewold, T.A., & Berckmans, D. (2015). Discerning pig screams in production environments. PloS one, 10(4), e0123111. 10.1371/journal.pone.0123111

37. Villain, A. S., Lanthony, M., Guérin, C., & Tallet, C. (2020a). Manipulable object and human contact: preference and modulation of emotional states in weaned pigs. Frontiers in Veterinary Science, 7, 577433. 10.3389/fvets.2020.577433

38. Villain, A. S., Hazard, A., Danglot, M., Guérin, C., Boissy, A., & Tallet, C. (2020b). Piglets vocally express the anticipation of pseudo-social contexts in their grunts. Scientific Reports, 10(1), 18496. 10.1038/s41598-020-75378-x

39. Villain, A. S., & Briefer, E. F. (2026). Vocal signals. In G. J. Mason, B. L. Nielsen, & M. T. Mendl (Eds.), Assessing Animal Welfare: A Guide to the Valid Use of Indicators of Affective States (pp. 80–102). John Wiley & Sons Ltd. 10.1002/9781394182237.ch4

40. Whitham, J. C., & Miller, L. J. (2024). Utilizing vocalizations to gain insight into the affective states of non-human mammals. Frontiers in veterinary science, 11, 1366933. 10.3389/fvets.2024.1366933

41. Winters, C. (2026). Tactile stimulation as a biomarker for positive animal welfare assessment: a systematic evaluation of current evidence and future directions. Zenodo. 10.5281/zenodo.20632728

