## Supporting information for "Pig vocalizations contain shared acoustic structure for humans and machines, but limited correspondence with presumed affective valence"

**This PDF file includes:**

Supporting text
Figures S1 to S18
Tables S1 to S7
Legends for Datasets S1 to S2
SI References

**Other supporting materials for this manuscript include the following:**

Datasets S1 to S2
Software S1

**Supporting Information Text**

**Methods**

**Replay-count analysis**

Replay counts were analyzed as an auxiliary measure of task difficulty and listener engagement. Because replay counts were non-negative count data with overdispersion, they were modeled using negative-binomial generalized linear mixed models with a log link. Separate models were fitted for Wave 1, Wave 2 context classification, and Wave 2 valence rating. Fixed effects included recording context, self-declared pig expertise group, sound-analysis experience, age group, gender, and acoustic characteristics of the vocalization: duration, loudness, and signal-to-noise ratio. Acoustic predictors were standardized before model fitting. Crossed random intercepts for participant and vocalization were included to account for repeated observations from the same participants and repeated presentation of the same vocalizations across participants. Model coefficients are reported as incidence-rate ratios (IRR) or as percentage changes in expected replay count. Because the Wave 2 valence task followed the context-classification task and used the same vocalizations, replay counts from the two Wave 2 tasks were analyzed separately and interpreted cautiously with respect to task difficulty.

**Classification-accuracy models**

Classification accuracy in Wave 2 was analyzed using binomial generalized linear mixed models with a logit link. Separate models were fitted for context-classification accuracy and directional valence-rating accuracy. For the context task, responses were coded as correct when the selected context matched the recording context of the vocalization. “No idea” responses were excluded from this model. For the valence task, responses were coded as correct when the selected directional valence matched the presumed context-derived valence of the vocalization. “No idea” responses and neutral responses were excluded from the directional valence-accuracy model.

Fixed effects included recording context, self-declared pig expertise group, gender, age group, professional-background category, sound-analysis experience, vocalization duration, loudness, and signal-to-noise ratio. Acoustic predictors were standardized before model fitting. Crossed random intercepts for participant and vocalization were included to account for repeated classifications by the same participants and repeated classification of the same vocalizations across participants. Model coefficients were exponentiated and are reported as odds ratios (ORs), where ORs above 1 indicate higher odds of correct classification and ORs below 1 indicate lower odds of correct classification.

**Results**

**Descriptive statistics and generalized mixed modeling replay counts**

In Wave 1, the survey was completed by 224 participants (70 males, 149 females, 5 other), who classified 8,960 vocalization presentations, representing 2,192 unique vocalizations, into 1,273 unique participant-defined categories. The survey in Wave 1 was initiated by 448 participants, indicating that 50.0% of respondents finished the survey task after starting it. Self-declared expertise with pigs was broadly distributed, with 88 participants classified as having low pig expertise (0–2), 75 as having medium expertise (3–6), and 63 as having high expertise (7–10). On average, participants played a vocalization 4.5 times (SD = 4.6), with a maximum of 66 plays.

Replay counts in Wave 1 were analyzed using a negative-binomial generalized linear mixed model with crossed random intercepts for participant and vocalization. The random-effect variance was substantially larger for participant identity than for vocalization identity, indicating that replay behavior varied more strongly between participants than between individual sounds after accounting for the fixed effects. In this model, pig expertise did not significantly predict replay count. Compared with participants with low pig expertise, participants with medium pig expertise had an estimated 11.1% higher replay count (IRR = 1.11, p = 0.267), whereas participants with high pig expertise had an estimated 6.7% lower replay count (IRR = 0.93, p = 0.496). Sound-analysis experience, age group, and gender showed no consistent association with replay count. Acoustic properties significantly predicted replay behavior: louder vocalizations were replayed less often (IRR = 0.95 per 1-SD increase in LUFS, $p = 4.36 \times 10^{-6}$), and longer vocalizations were replayed less often (IRR = 0.89 per 1-SD increase in duration, $p < 2 \times 10^{-16}$). Signal-to-noise ratio was not significantly associated with replay count (IRR = 0.99, p = 0.363). Several recording contexts also differed from the reference context (after nursing), with lower replay counts for barren, castration, fighting, novel-object, positive-conditioning, and restraint vocalizations.

In Wave 2, the survey was completed by 159 participants (53 males, 98 females, 8 other), who classified 3,975 vocalization presentations, representing 1,357 unique vocalizations, using predefined categories for context (18 levels and “no idea”) and valence (5 levels and “no idea”). The survey in Wave 2 was initiated by 374 participants, indicating that 42.5% of respondents finished the survey task after starting it. Self-declared expertise with pigs showed a distribution weighted toward lower expertise, with 75 participants classified as having low pig expertise (0–2), 42 as having medium expertise (3–6), and 44 as having high expertise (7–10). On average, participants played a vocalization 4.5 times (SD = 4.5), with a maximum of 51 plays, during the context-classification task. During the valence-rating task, participants played a vocalization 3.1 times on average (SD = 2.8), with a maximum of 31 plays.

Replay counts in Wave 2 were analyzed separately for the context- and valence-rating tasks using negative-binomial generalized linear mixed models with crossed random intercepts for participant and vocalization. In the context-classification task, replay behavior again varied primarily between participants, whereas the estimated random-effect variance for vocalization identity was negligible after accounting for the fixed effects. Compared with participants with low pig expertise, participants with medium pig expertise replayed vocalizations more often (IRR = 1.31, p = 0.044), while the corresponding estimate for participants with high pig expertise was positive but did not reach significance (IRR = 1.27, p = 0.073). Acoustic properties also predicted replay count: louder vocalizations were replayed less often (IRR = 0.95 per 1-SD increase in LUFS, $p = 8.46 \times 10^{-5}$), and longer vocalizations were replayed less often (IRR = 0.84 per 1-SD increase in duration, $p < 2 \times 10^{-16}$). Signal-to-noise ratio was not significantly associated with replay count (IRR = 1.01, p = 0.443). Relative to the reference context (after nursing), replay counts were lower for castration and novel-object vocalizations and higher for positive-conditioning vocalizations.

In the Wave 2 valence-rating task, pig expertise did not significantly predict replay count. Compared with participants with low pig expertise, replay counts did not differ significantly for participants with medium pig expertise (IRR = 1.08, p = 0.487) or high pig expertise (IRR = 1.03, p = 0.763). Sound-analysis experience and gender were not significantly associated with replay count. Older age groups showed lower replay counts for participants aged 65–74 and 75–84 compared with the reference age group, although these age effects should be interpreted cautiously because of likely small group sizes. As in the other replay models, acoustic properties predicted replay behavior: louder vocalizations were replayed less often (IRR = 0.93 per 1-SD increase in LUFS, $p = 3.08 \times 10^{-6}$), and longer vocalizations were replayed less often (IRR = 0.88 per 1-SD increase in duration, $p = 2.86 \times 10^{-11}$). Signal-to-noise ratio was not significantly associated with replay count (IRR = 1.00, p = 0.799). Several contexts had lower replay counts than the reference context, including castration, crushing, fighting, negative conditioning, novel object, and restraint.

Together, these replay-count models indicate that replay behavior was influenced more consistently by general participant-level replay tendencies and basic acoustic properties than by pig-related expertise. Across waves and tasks, louder and longer vocalizations were replayed less often, whereas signal-to-noise ratio showed no consistent effect. Pig expertise was not associated with replay count in Wave 1 or in the Wave 2 valence task, and showed only limited evidence of an association in the Wave 2 context-classification task.

**Classification-accuracy mixed models**

Classification accuracy in Wave 2 was analyzed using binomial generalized linear mixed models with a logit link. Models were fitted separately for context-classification accuracy and directional valence-rating accuracy. Coefficients were exponentiated and are reported as odds ratios (ORs), where values above 1 indicate higher odds of a correct classification and values below 1 indicate lower odds of a correct classification.

For context classification, there was no evidence that pig expertise significantly predicted classification accuracy. Compared with participants with low pig expertise, the odds of correct context classification did not differ significantly for participants with medium pig expertise (OR = 1.06, p = 0.722) or high pig expertise (OR = 0.70, p = 0.237). Gender, age group, professional background, and sound-analysis experience also showed no clear associations with context-classification accuracy. The only acoustic predictor significantly associated with context accuracy was duration. Longer vocalizations had higher odds of being classified correctly (OR = 1.35 per 1-SD increase in duration, $p = 9.93 \times 10^{-5}$). Loudness (OR = 1.08, p = 0.411) and signal-to-noise ratio (OR = 1.03, p = 0.714) were not significantly associated with context accuracy. Among recording contexts, only long isolation differed significantly from the reference context (after nursing), showing lower odds of correct context classification (OR = 0.26, p = 0.027). The participant-level random-effect variance was estimated as zero, whereas vocalization identity showed a small random-effect variance, indicating little residual participant-level variation in context-classification accuracy after accounting for the fixed effects.

For directional valence rating, recording context and acoustic properties were more strongly associated with accuracy. Pig expertise again did not significantly predict accuracy. Compared with participants with low pig expertise, participants with medium pig expertise showed a non-significant tendency toward higher odds of correct directional valence rating (OR = 1.23, p = 0.085), whereas participants with high pig expertise did not differ from the low-expertise group (OR = 1.05, p = 0.802). Gender, professional background, and sound-analysis experience showed no clear associations with directional valence accuracy. Participants aged 65–74 had lower odds of correct directional valence rating than the reference age group of 18-25 years (OR = 0.45, p = 0.045), although these age effects should be interpreted cautiously because of small group sizes.

Directional valence accuracy varied strongly by recording context. Relative to the reference context (after nursing), the odds of correct directional valence rating were higher for castration (OR = 5.71, $p = 2.73 \times 10^{-6}$), crushing (OR = 1.97, p = 0.024), fighting (OR = 5.13, $p = 6.25 \times 10^{-7}$), positive conditioning (OR = 2.35, p = 0.041), and restraint (OR = 7.35, $p = 4.20 \times 10^{-7}$). In contrast, the odds of correct directional valence rating were lower for barren (OR = 0.10, $p = 1.18 \times 10^{-7}$), negative conditioning (OR = 0.16, p = 0.0002), run (OR = 0.50, p = 0.023), short isolation (OR = 0.44, p = 0.006), and surprise vocalizations (OR = 0.34, p = 0.004). Longer vocalizations had higher odds of correct directional valence rating (OR = 1.42 per 1-SD increase in duration, $p = 1.91 \times 10^{-5}$), and louder vocalizations also had slightly higher odds of correct directional valence rating (OR = 1.16 per 1-SD increase in LUFS, p = 0.048). Signal-to-noise ratio was not significantly associated with valence accuracy (OR = 1.02, p = 0.750). The random-effect variance for participant identity was again estimated as zero, whereas vocalization identity showed substantial residual variance, indicating that accuracy depended more on which vocalization was classified than on stable participant-level differences after accounting for the fixed effects.

Together, these accuracy models indicate that participant-related predictors, including pig expertise, explained little variation in classification accuracy once recording context, acoustic properties, and repeated observations were taken into account. Context-classification accuracy was weakly structured, with longer sounds being more likely to be classified correctly. Directional valence accuracy showed stronger context dependence, with higher accuracy for several highly aversive contexts such as castration, fighting, restraint, and crushing. This supports the conclusion that valence rating was more structured than exact context classification, but that this structure was uneven across contexts and not primarily explained by pig-related expertise.

**Figures**

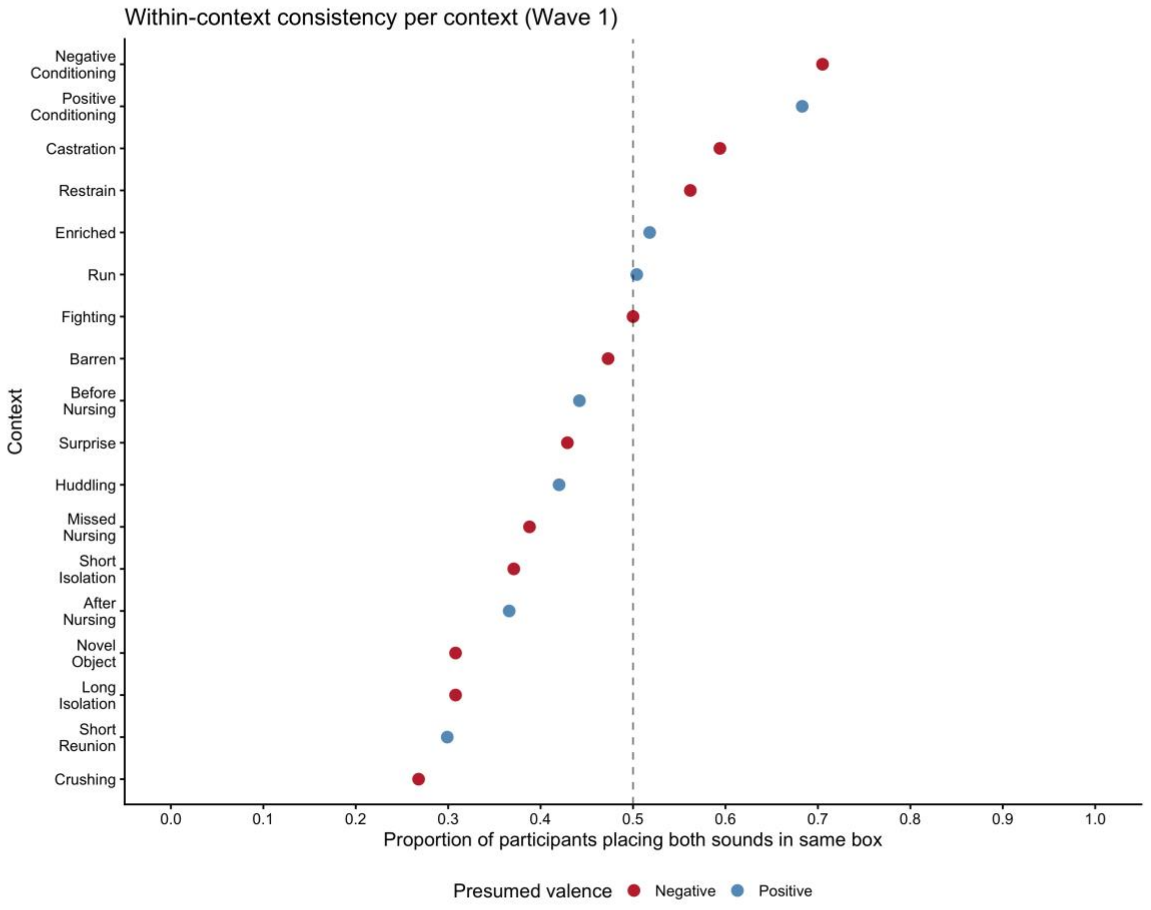


**Fig. S1. Within-context consistency of Wave 1 free classification.** Points show, for each recording context, the proportion of participants who placed both sounds from the same context in the same participant-defined category. The dashed vertical line indicates 50% consistency. Points are colored by presumed valence as defined in Briefer et al. (2022a). The highest within-context consistency was observed for negative conditioning (70.5%), positive conditioning (68.3%), and castration (59.4%), whereas the lowest consistency was observed for crushing (26.8%), short reunion (29.7%), and novel object test (30.8%).

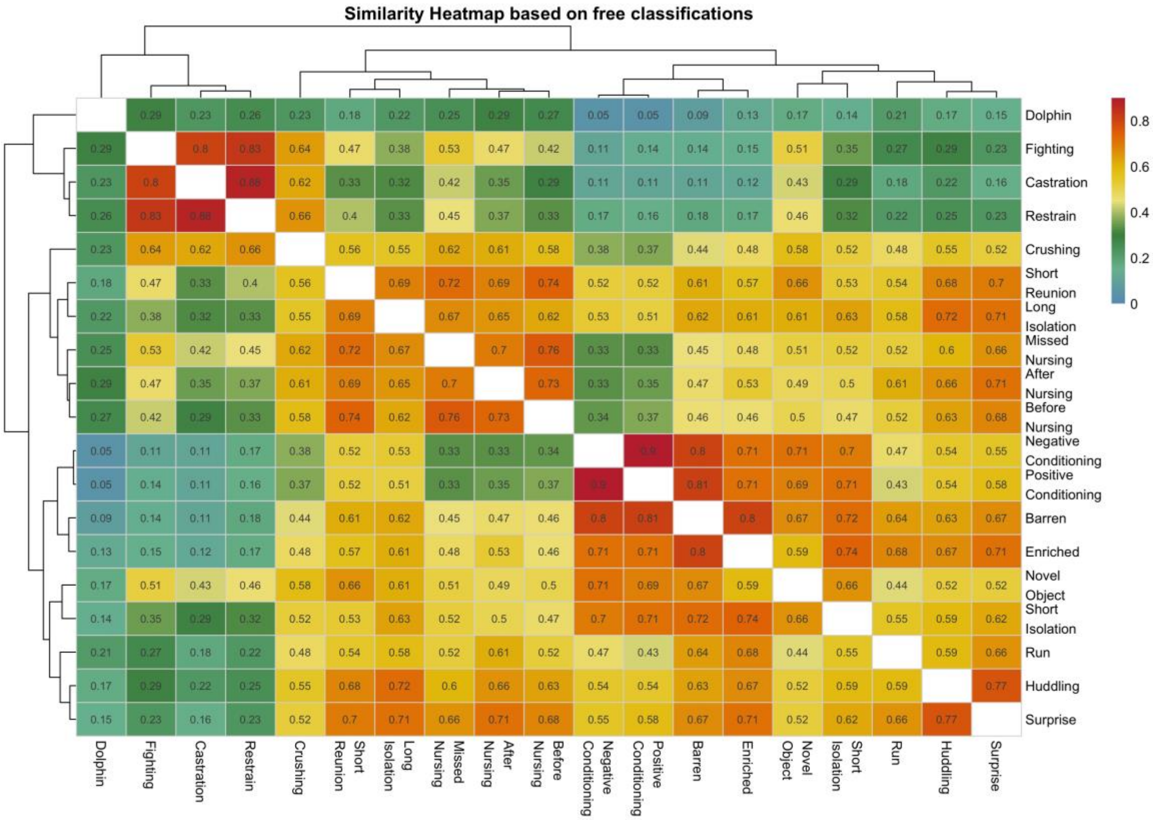


**Fig. S2. Human-derived clustering similarity among pig vocalization contexts based on free classification.** Heatmap showing context-level similarity derived from participant free-classification groupings in Wave 1 (N = 224 participants). Similarity values represent the proportion of participants who assigned vocalizations from each pair of recording contexts to the same freely chosen category. Higher values indicate greater perceived similarity. Rows and columns are ordered according to the context clusters derived from the classification structure; the diagonal is omitted.

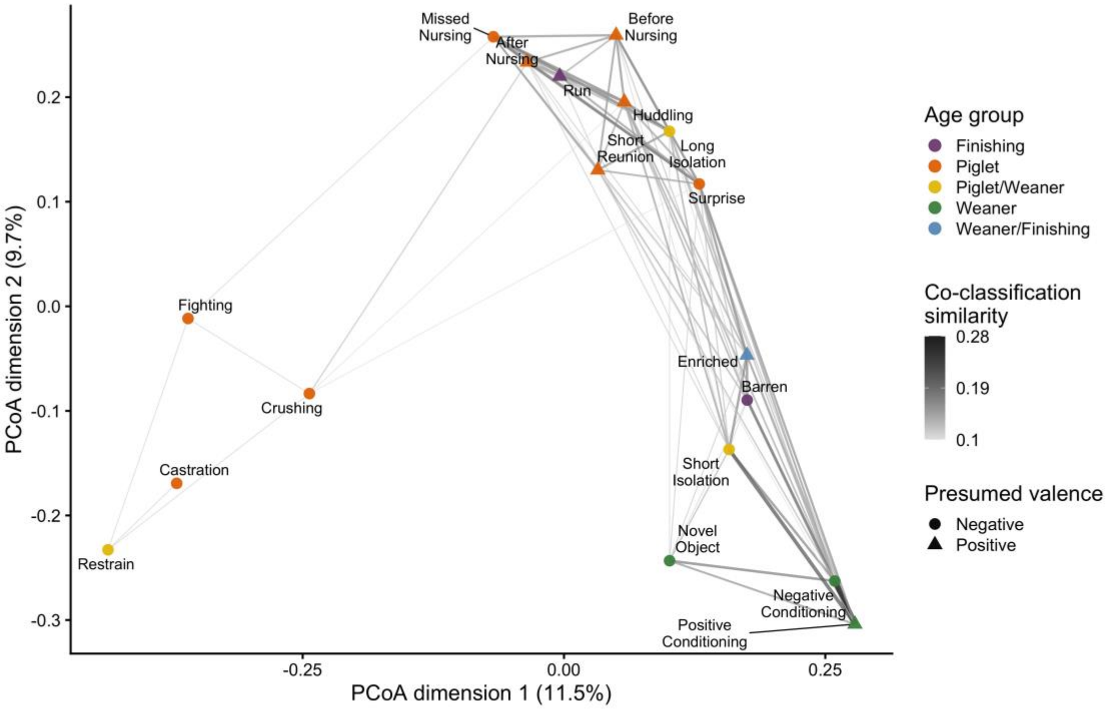


**Fig. S3. Human perceptual similarity among pig vocalization contexts based on forced-choice classification on context (N = 159 participants; Wave 2).** Two-dimensional principal coordinates analysis of human forced-choice context classification similarity among pig vocalization contexts. Points represent contexts; distances reflect dissimilarity in participant co-classification patterns. Lines connect context pairs with similarity ≥ 0.10, with thicker lines indicating stronger similarity. As there were 18 pre-defined context categories for 25 vocalizations per participant, co-classification similarities are much lower compared to other classification tasks. Point color indicates age group and point shape indicates presumed valence as in Briefer et al., 2022a.

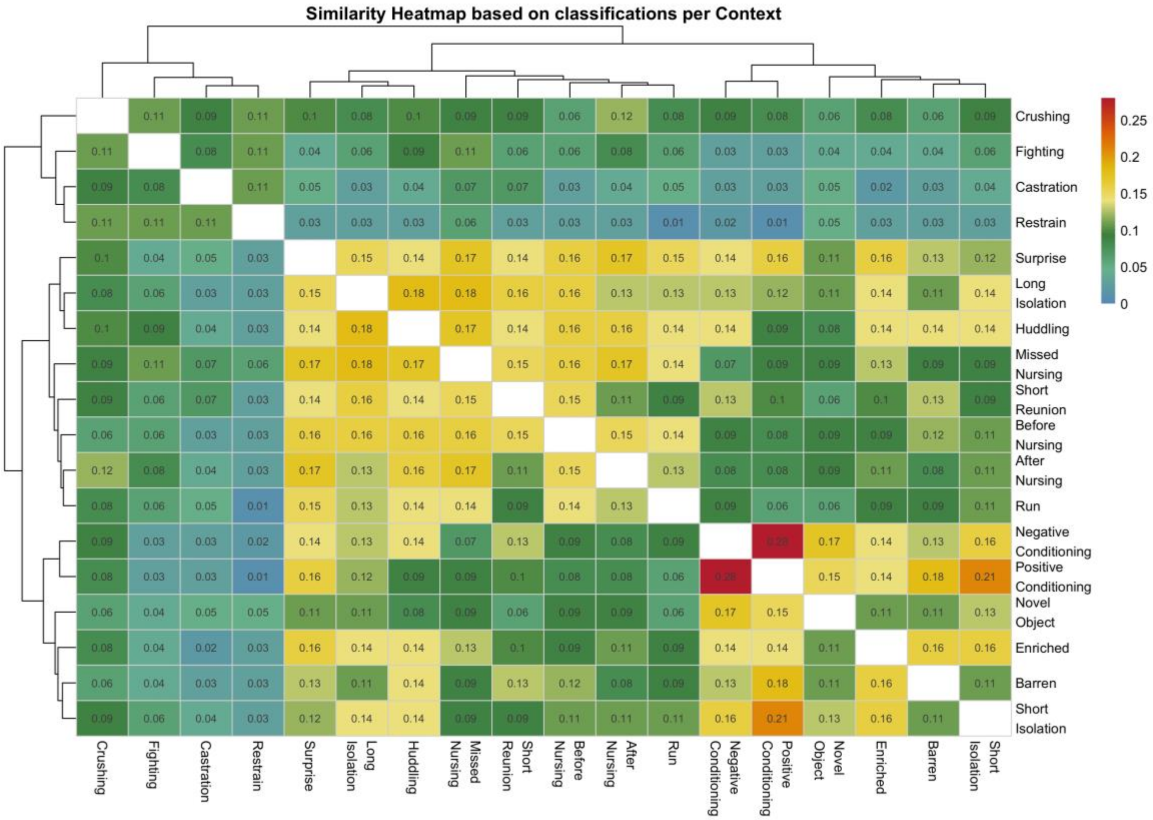


**Fig. S4. Human-derived similarity among pig vocalization contexts based on forced-choice context classification.** The heatmap showing context-level similarity derived from participant context classifications in Wave 2 (N = 159 participants). Similarity values represent the proportion of participants who assigned vocalizations from each pair of recording contexts to the same context. Higher values indicate greater perceived similarity in context classification. As there were 18 pre-defined context categories for 25 vocalizations per participant, co-classification similarities are much lower compared to other classification tasks. Rows and columns are ordered according to the context clusters derived from the classification structure; the diagonal is omitted.

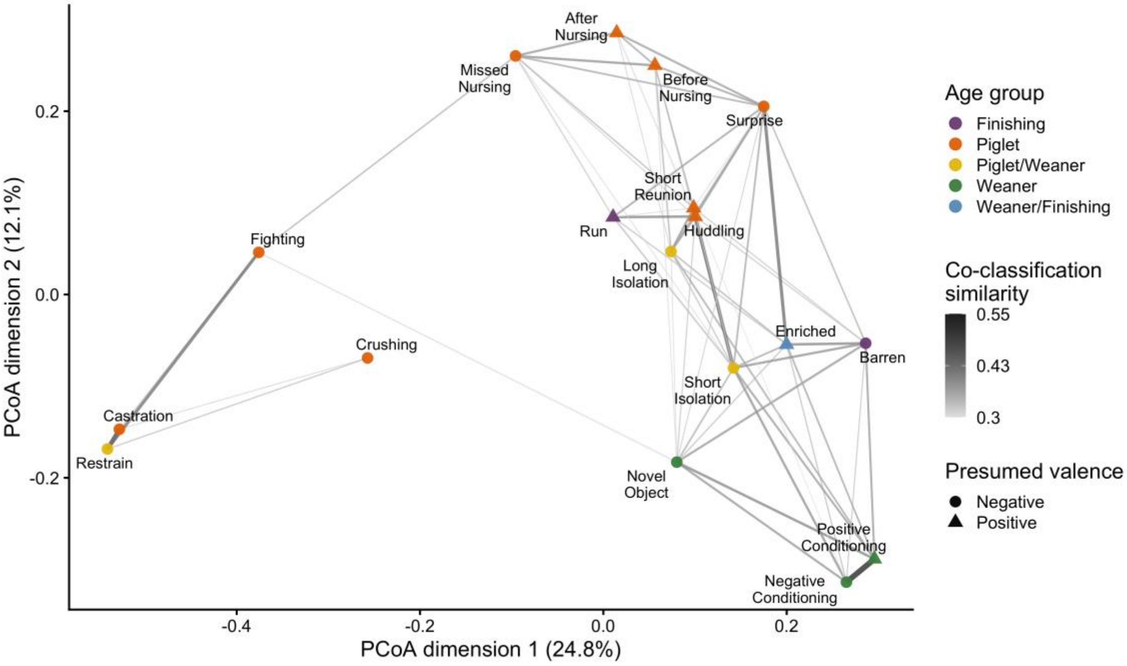


**Fig. S5. Human perceptual similarity among pig vocalization contexts based on forced-choice rating of valence (N = 159 participants; Wave 2).** Two-dimensional principal coordinates analysis of human forced-choice valence rating similarity among pig vocalization contexts. Points represent contexts; distances reflect dissimilarity in participant co-classification patterns. Lines connect context pairs with similarity ≥ 0.10, with thicker lines indicating stronger similarity. Point color indicates age group and point shape indicates presumed valence as in Briefer et al., 2022a.

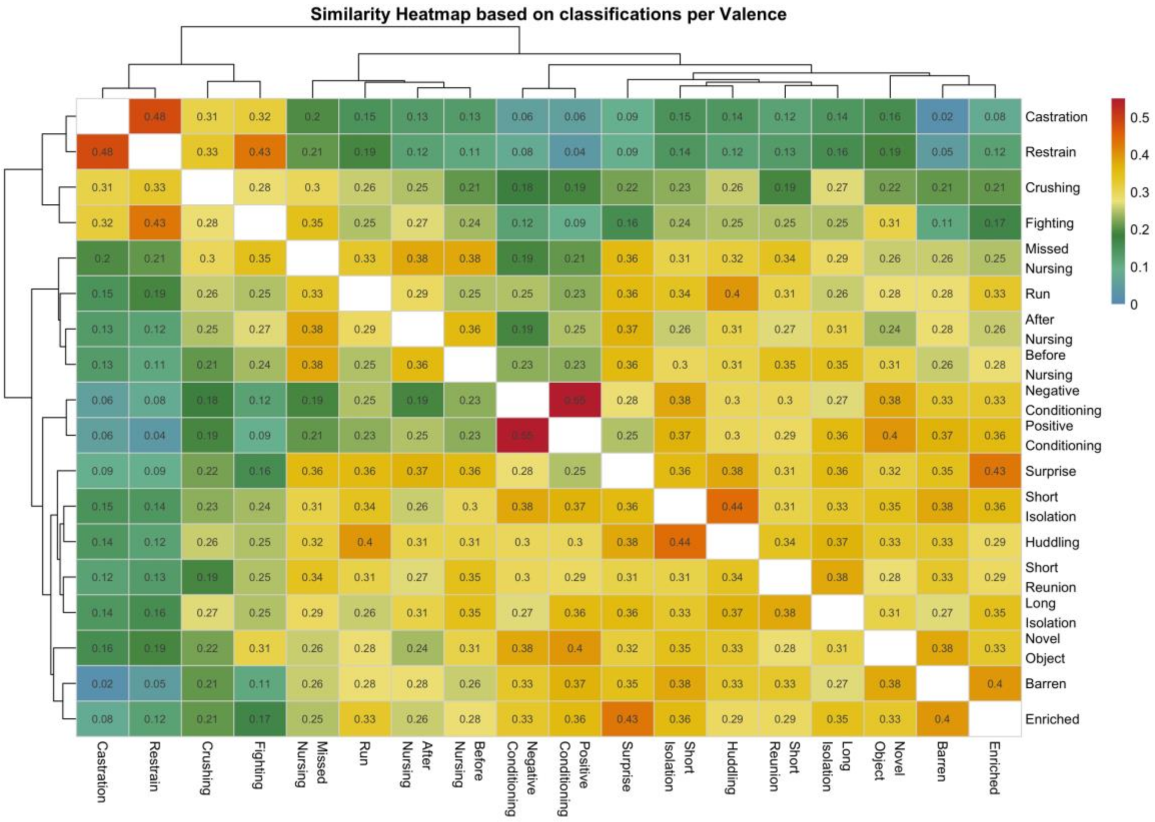


**Fig. S6. Human-derived similarity among pig vocalization contexts based on forced-choice valence rating.** Heatmap showing context-level similarity derived from participant valence rating in Wave 2 (N = 159 participants). Similarity values represent the proportion of participants who assigned vocalizations from each pair of recording contexts to the same valence score. Higher values indicate greater perceived similarity in valence rating. Rows and columns are ordered according to the context clusters derived from the classification structure; the diagonal is omitted.

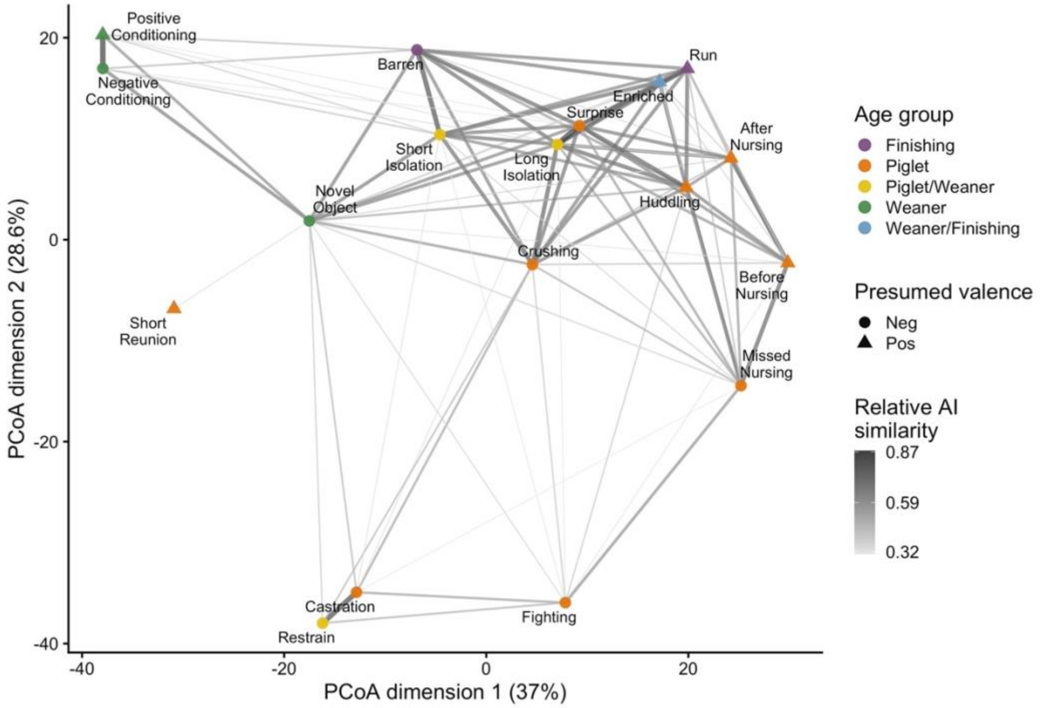


**Fig. S7. Machine-learned acoustic similarity among pig vocalization contexts for the context-optimized model.** Two-dimensional principal coordinates analysis of the AI-derived context-distance matrix based on scaled ResNet50 embedding centroids. Points represent recording contexts; distances reflect Euclidean dissimilarity between context-level embedding centroids. Lines connect context pairs above the prespecified relative AI-similarity threshold, with thicker and darker lines indicating smaller Euclidean distance, and therefore greater relative AI similarity. Point color indicates age group and point shape indicates presumed valence as in Briefer et al., 2022a.

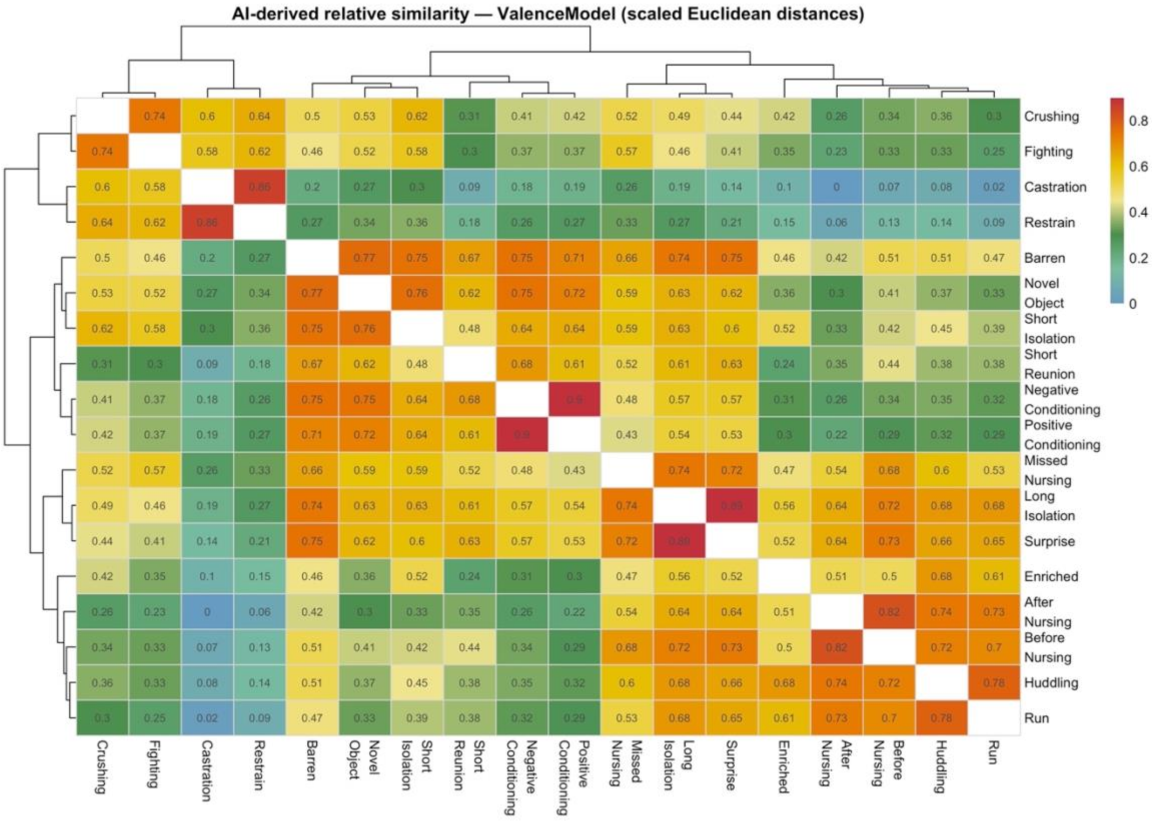


**Fig. S8. AI-derived relative similarity among pig vocalization contexts optimized for context.** Heatmap showing relative similarity between recording contexts based on scaled ResNet50 embedding centroids. Euclidean distances between context centroids were normalized by the maximum pairwise distance and converted to relative similarity as 1 − normalized distance. Higher values indicate smaller embedding distances and therefore greater machine-learned acoustic similarity. Rows and columns are clustered using the AI-derived distance matrix; the diagonal is omitted.

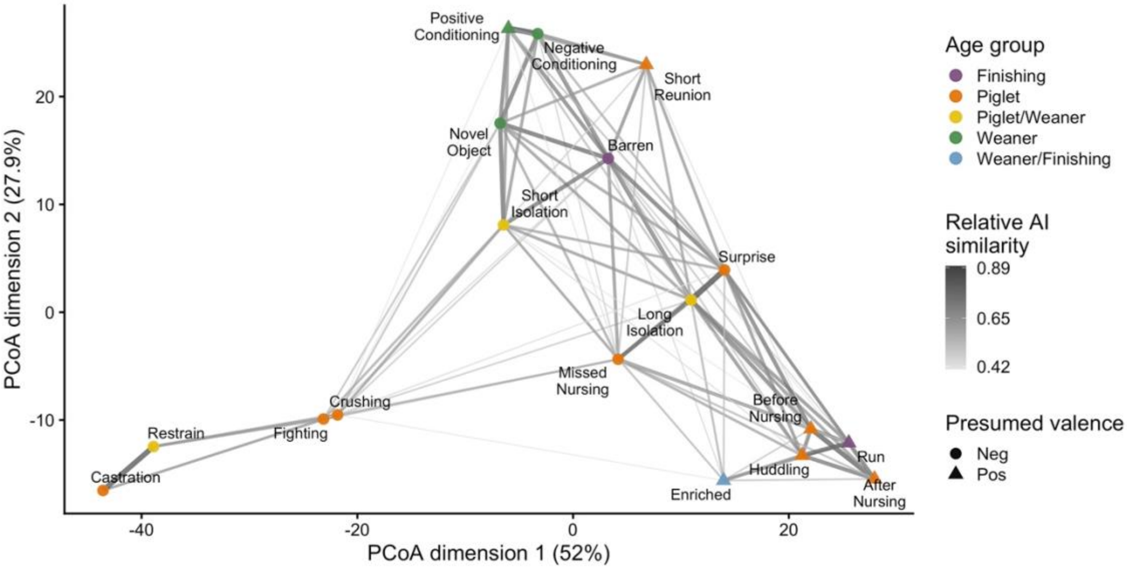


**Fig. S9. Machine-learned acoustic similarity among pig vocalization contexts for the valence-optimized model.** Two-dimensional principal coordinates analysis of the AI-derived context-distance matrix based on scaled ResNet50 embedding centroids. Points represent recording contexts; distances reflect Euclidean dissimilarity between context-level embedding centroids. Lines connect context pairs above the prespecified relative AI-similarity threshold, with thicker and darker lines indicating smaller Euclidean distance, and therefore greater relative AI similarity. Point color indicates age group and point shape indicates presumed valence as in Briefer et al., 2022a.

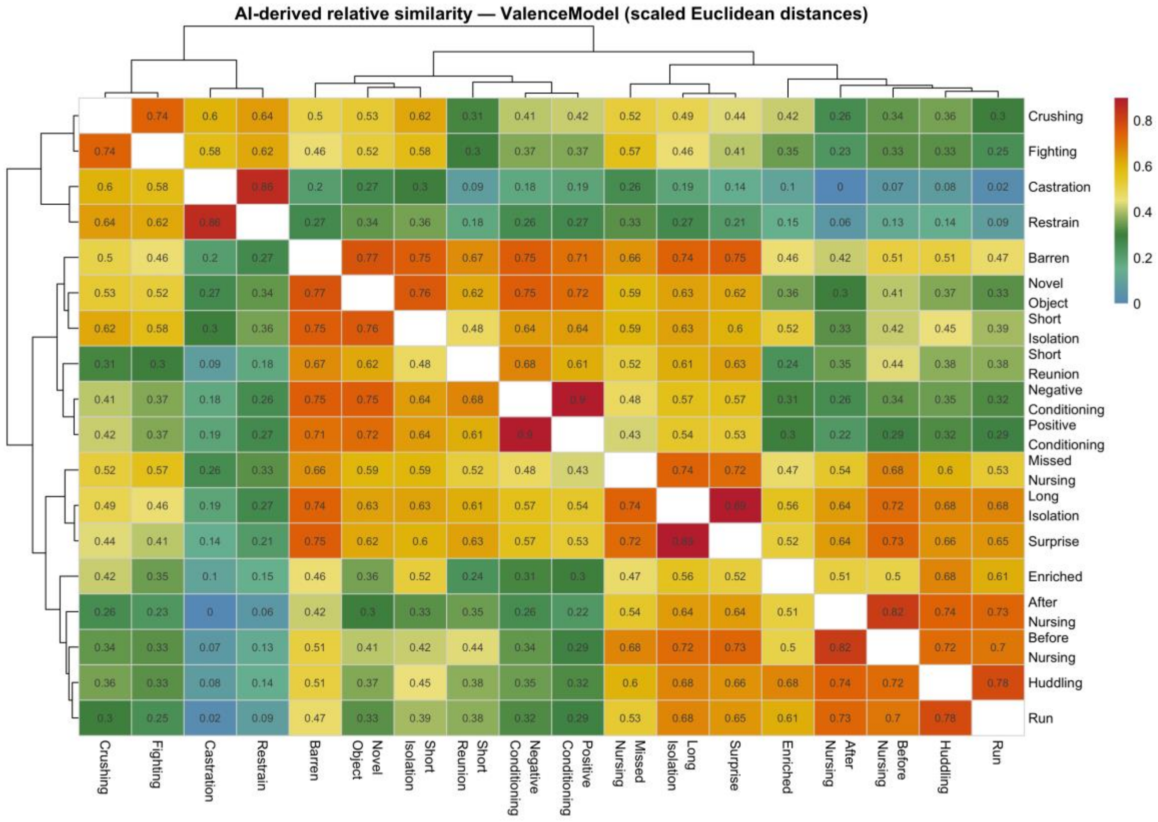


**Fig. S10. AI-derived relative similarity among pig vocalization contexts optimized for valence.** Heatmap showing relative similarity between recording contexts based on scaled ResNet50 embedding centroids. Euclidean distances between context centroids were normalized by the maximum pairwise distance and converted to relative similarity as 1 − normalized distance. Higher values indicate smaller embedding distances and therefore greater machine-learned acoustic similarity. Rows and columns are clustered using the AI-derived distance matrix; the diagonal is omitted.

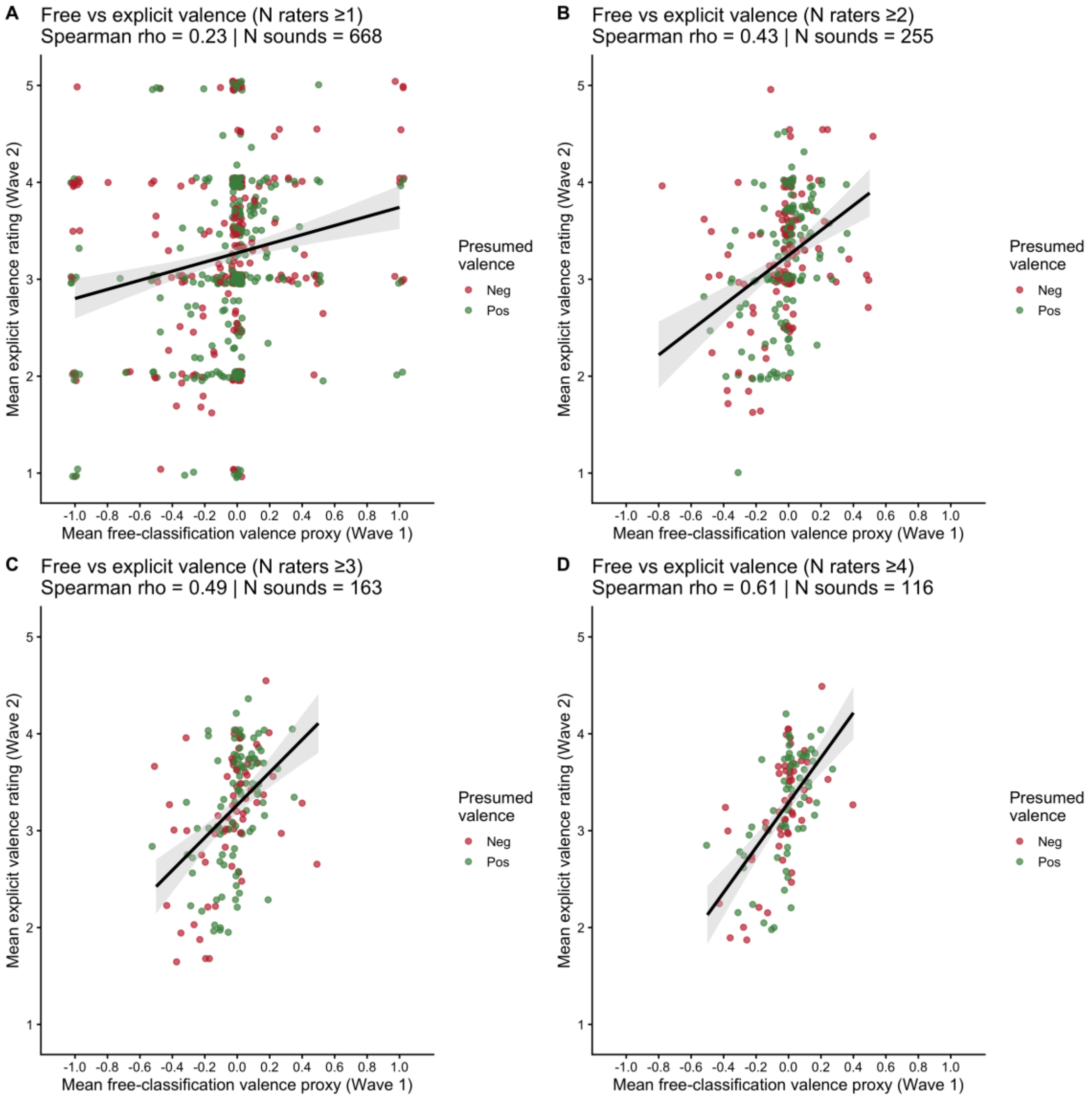


**Fig. S11. Free-classification labels still contain a repeatable valence signal compared to forced-choice valence scoring after excluding high-intensity aversive contexts (castration, restraint, fighting, and crushing).** Relationship between a valence score proxy derived from Wave 1 free-classification labels and explicit Wave 2 valence ratings for the same vocalizations. Feeling- and emotion-related free labels were converted to a signed proxy score, with positive labels coded as 1, negative labels as −1, and mixed labels or labels without feeling or emotion labels scored as 0. Explicit Wave 2 valence ratings were scored on a five-point Likert scale from very negative to very positive. Each point represents the average values for one sound over participants. Points were jittered to show overlapping points. Panels show increasingly strict inclusion thresholds based on the minimum number of raters per sound in both waves: at least one rater per wave (A), at least two raters per wave (B), at least three raters per wave (C), and at least four raters per wave (D). Blue lines show linear fits with 95% confidence intervals. Spearman's ρ and the number of sounds included are shown within each panel.

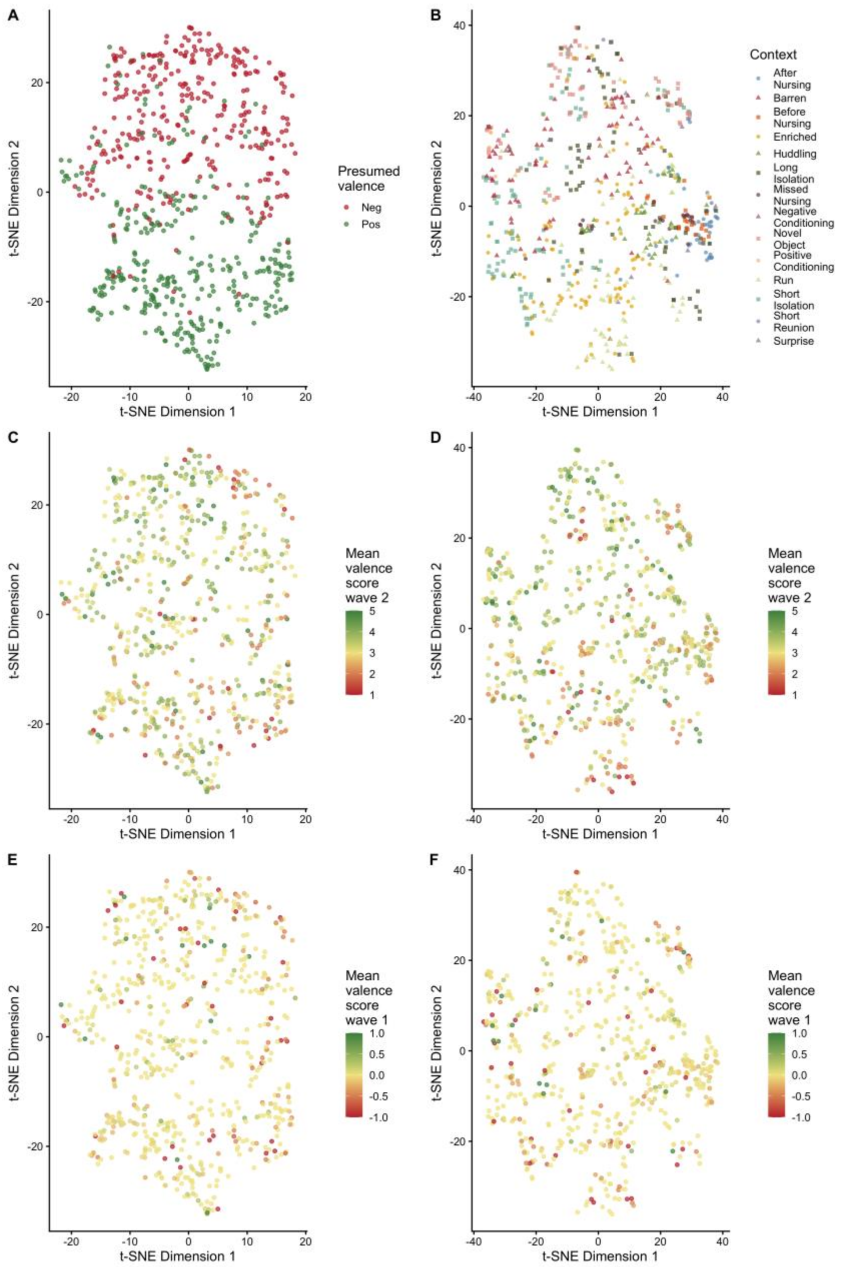


**Figure S12. Low-dimensional projections of convolutional neural network embeddings for pig vocalizations, excluding the high-intensity aversive contexts (castration, restraint, fighting and crushing).** t-distributed stochastic neighbor embedding (t-SNE) projections of ResNet50 embedding vectors from models fine-tuned for presumed valence (A, C, E) or context classification (B, D, F). Each point represents one vocalization and only the 668 vocalizations that were scored in wave 1 and wave 2 are shown. Points are colored by presumed valence in A as per Briefer et al. 2022a, by recording context in B, by mean explicit wave 2 valence rating in C and D and by mean inferred wave 1 valence rating in E and F.

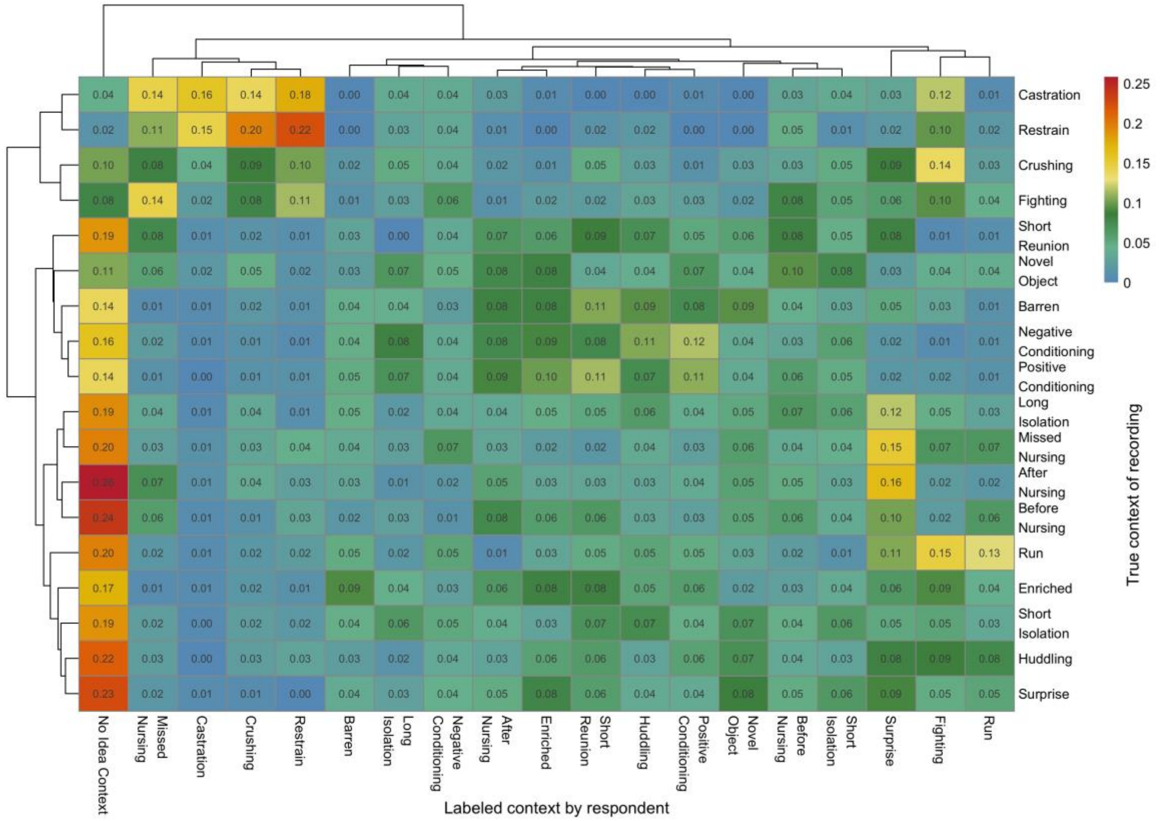


**Fig. S13. Confusion matrix of forced-choice context labeling in wave 2.** Heatmap showing the proportion of vocalizations from each true recording context assigned to each response category by participants. Rows indicate the true context of the recording, and columns indicate the context label selected by the respondent. Values are row-normalized, so each row shows the distribution of participant responses for a given true context. Rows and columns are hierarchically clustered to visualize similarities in classification patterns. Higher values indicate more frequent assignments to that response category. The “No Idea Context” column represents trials for which participants did not assign a context label.

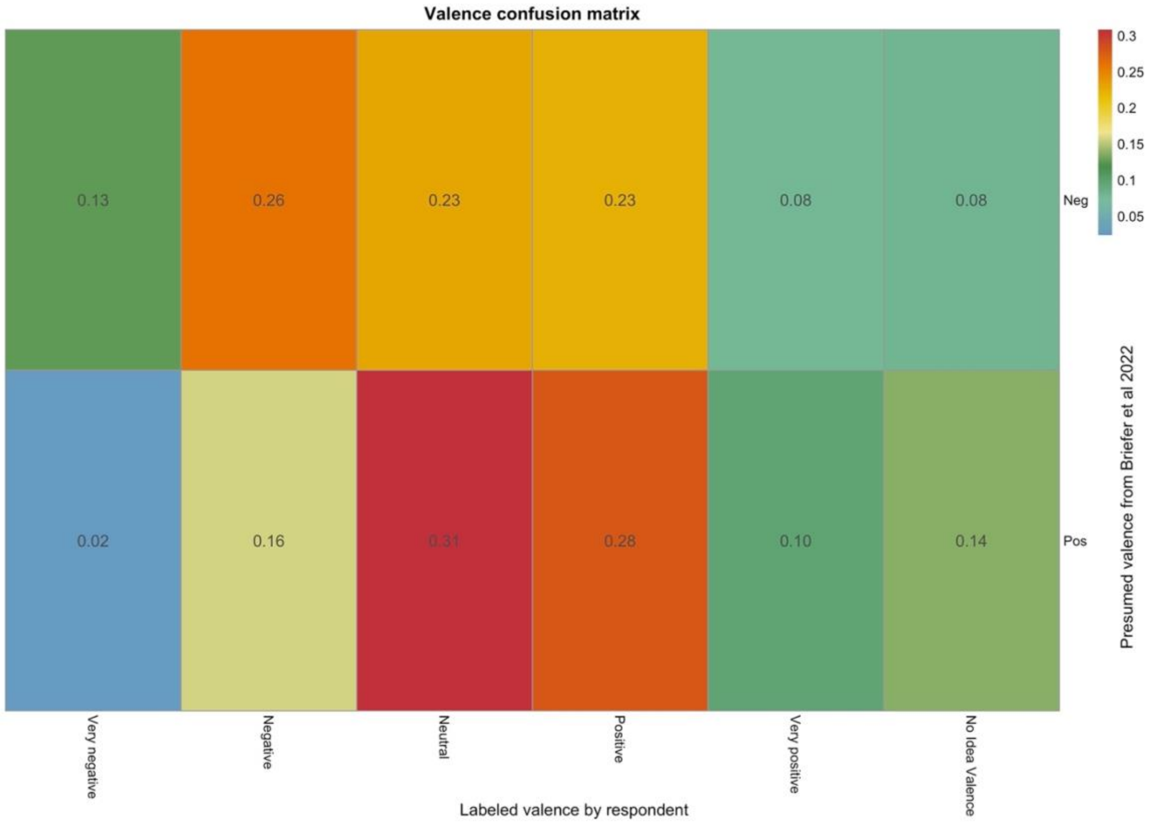


**Fig. S14. Confusion matrix of forced-choice valence labeling in wave 2.** Heatmap showing the proportion of source-negative and source-positive vocalizations assigned to each valence response category by participants. Rows indicate presumed valence as defined in Briefer et al., 2022a, and columns indicate the valence label selected by the respondent on the five-point ordinal scale. Values are row-normalized, so each row shows the distribution of participant responses for a given presumed valence class. Higher values indicate more frequent assignments to that response category. “No Idea Valence” responses were excluded from this heatmap.

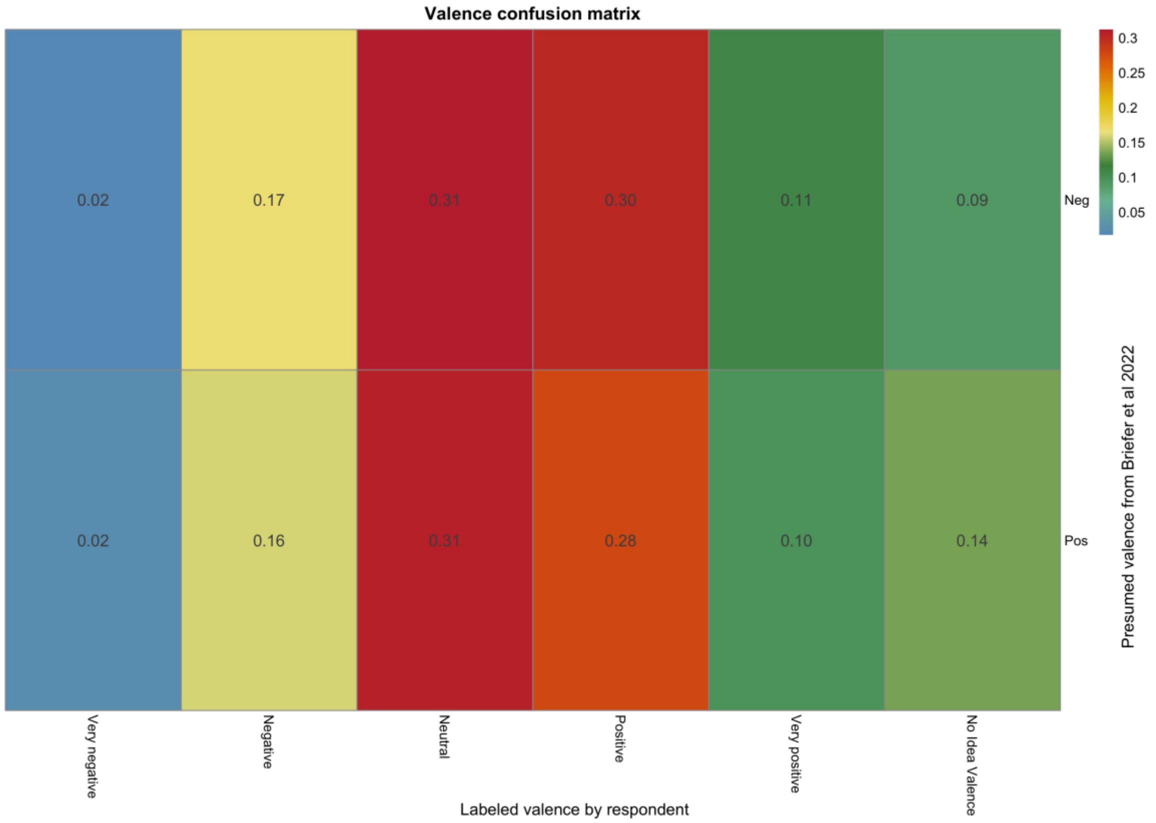


**Fig. S15. Confusion matrix of forced-choice valence labeling in wave 2 excluding the high-intensity aversive contexts (castration, restraint, fighting and crushing).** Heatmap showing the proportion of source-negative and source-positive vocalizations assigned to each valence response category by participants. Rows indicate presumed valence as defined in Briefer et al., 2022a, and columns indicate the valence label selected by the respondent on the five-point ordinal scale. Values are row-normalized, so each row shows the distribution of participant responses for a given presumed valence class. Higher values indicate more frequent assignments to that response category. “No Idea Valence” responses were excluded from this heatmap. After excluding the high-intensity aversive contexts (castration, restraint, fighting and crushing), the classification accuracy was not significantly different anymore from random.

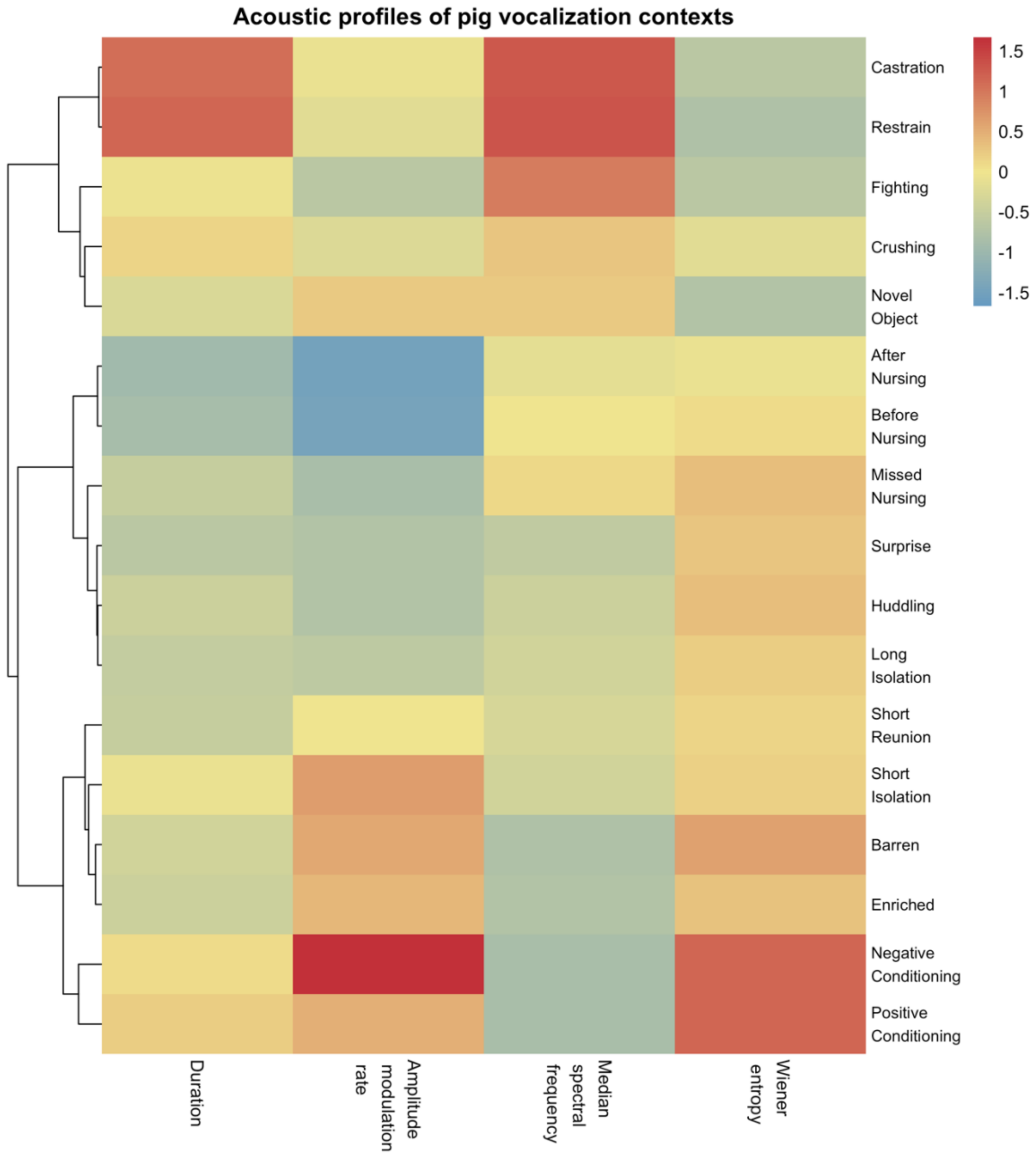


**Fig. S16. Context-level acoustic profiles of pig vocalizations based on four conventional acoustic measures.** Heatmap showing mean standardized values of call duration, amplitude-modulation rate, median spectral frequency (Q50), and Wiener entropy for each recording context. Acoustic measures were standardized across individual vocalizations before averaging within context; warmer colors indicate values above the overall mean and cooler colors values below the overall mean. Contexts were hierarchically clustered using Euclidean distances between their four-dimensional acoustic profiles and Ward's D2 linkage. Columns were not clustered.

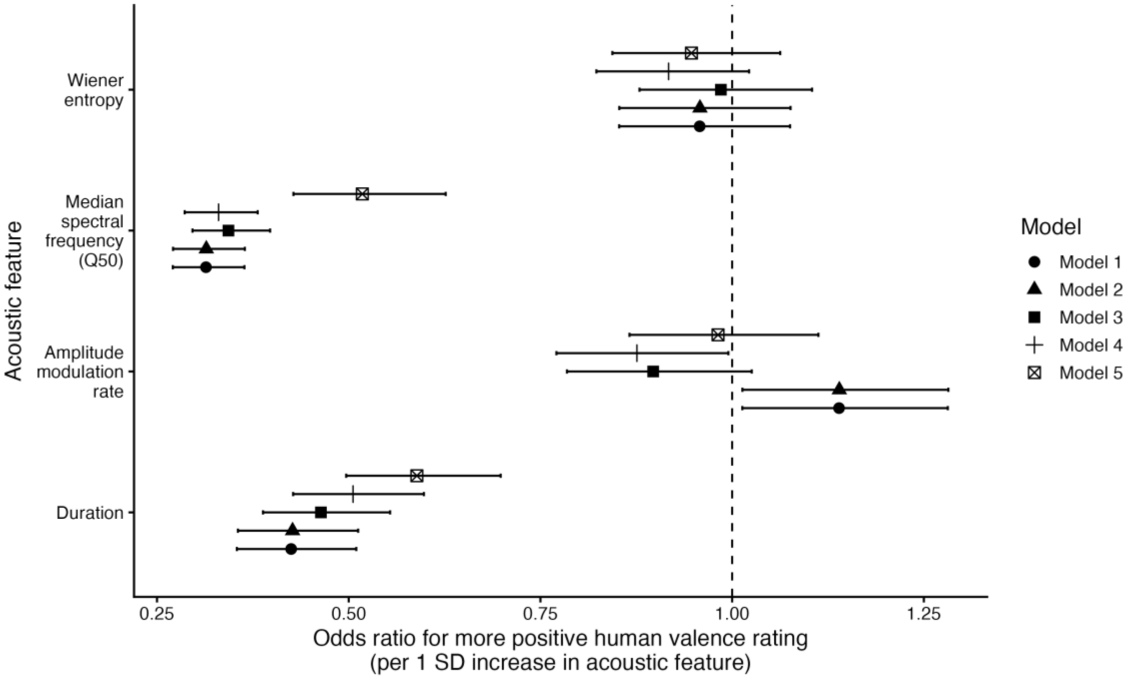


**Fig. S17. Acoustic predictors of human-perceived valence across sequentially adjusted models.** Odds ratios and 95% confidence intervals represent the association between a one-standard-deviation increase in each acoustic feature and the odds of assigning a more positive valence rating. Model 1 included the four acoustic predictors; Model 2 additionally included listener characteristics; Model 3 additionally included pig characteristics; Model 4 additionally included recording characteristics; and Model 5 additionally included recording context and call type. All models included random intercepts for participant and vocalization. The dashed line indicates an odds ratio of 1 (no association).

**Fig. S18. Participant-facing online survey interface.** Representative screenshots of the Project Oink online survey. (A) Information and consent form. (B, C) Wave 1 free-classification interface, in which participants listened to vocalizations and created participant-defined categories by dragging audio files into boxes. (D, F) Participant background and demographic questions. (E) Wave 2 forced-choice context-classification interface, including the context definitions and predefined context boxes. (G) Wave 2 forced-choice valence-rating interface, in which participants assigned vocalizations to five valence categories or to “No idea.” (H) Confirmation screen before proceeding to the next part of the survey. (I) Survey completion screen. Screenshots are shown to document the task interface and response options presented to participants.

### Tables

**Table S1. Detailed evidential basis for the inferential audit of pig vocalizations as indicators of affective state.** This table provides the study-level evidence underlying the summary classifications reported in Table 1. For each study and auxiliary assumption (A1 to A5), we document the relevant operationalization, why the assumption is required for the affective inference, how the study addressed it, the empirical or theoretical evidence available, the resulting evidential status, and the consequence for interpretation if the assumption does not hold. We additionally distinguish the literal empirical finding from the authors' affective interpretation and identify the inferential step required to move between them.

The audit separates evidence that vocalizations discriminate experimental or observational conditions from evidence that such differences specifically reflect emotional valence or another affective construct. Accordingly, lack of support for an auxiliary assumption does not imply that the proposed affective interpretation is false; rather, it indicates that the available study design or evidence does not uniquely establish that interpretation. Likewise, **Not tested** is distinguished from **Inconclusive** and from **Contrasted**, because absence of a test, unresolved evidence, and evidence against invariance represent different inferential situations.

Table S1 also records relevant sources of evidential dependence, including reuse of animals or vocalization datasets, inheritance of affective labels from previous studies, and reliance on previously proposed acoustic–affect mappings. These details are used to avoid treating reanalysis, pooled datasets, or inherited construct assumptions as fully independent tests of the same affective claim.

See Additional Excel file: “TableS1_Audit_Vocalization_affect.xlsx”

**Table S2. Feelings-related labels assigned to positive or negative valence in free classification.** The maximum amount of feelings labels was 448 per context category, as each of the 224 respondents got 2 vocalizations per context category.

| Context of vocalization | Number of vocalizations scored with feelings label | Proportion with feelings label | Relative proportion negatively labeled | Relative proportion positively labeled | Relative proportion mixed labeled |
|---|---|---|---|---|---|
| **Castration** | 161 | 0.359 | 0.957 | 0.025 | 0.018 |
| **Restrain** | 158 | 0.353 | 0.924 | 0.044 | 0.032 |
| **Fighting** | 133 | 0.297 | 0.865 | 0.083 | 0.052 |
| **Crushing** | 123 | 0.275 | 0.789 | 0.146 | 0.065 |
| **Novel Object** | 105 | 0.234 | 0.486 | 0.381 | 0.133 |
| **Missed Nursing** | 91 | 0.203 | 0.571 | 0.286 | 0.143 |
| **Barren** | 87 | 0.194 | 0.379 | 0.414 | 0.207 |
| **Short Isolation** | 85 | 0.19 | 0.6 | 0.247 | 0.153 |
| **Enriched** | 84 | 0.188 | 0.524 | 0.262 | 0.214 |
| **Positive Conditioning** | 83 | 0.185 | 0.289 | 0.434 | 0.277 |
| **Run** | 81 | 0.181 | 0.716 | 0.148 | 0.136 |
| **Long Isolation** | 80 | 0.179 | 0.512 | 0.312 | 0.176 |
| **Negative Conditioning** | 78 | 0.174 | 0.295 | 0.449 | 0.256 |
| **Short Reunion** | 74 | 0.165 | 0.338 | 0.473 | 0.189 |
| **Huddling** | 72 | 0.161 | 0.389 | 0.444 | 0.167 |
| **After Nursing** | 67 | 0.15 | 0.567 | 0.299 | 0.134 |
| **Before Nursing** | 66 | 0.147 | 0.455 | 0.409 | 0.136 |
| **Surprise** | 63 | 0.141 | 0.413 | 0.429 | 0.158 |
| **Dolphin** | 60 | 0.134 | 0.767 | 0.2 | 0.033 |

**Table S3. Context-level performance in the Wave 2 forced-choice context-classification task.** Values are calculated per true recording context after excluding duplicate trials. Recall is the proportion of sounds from a given context correctly assigned to that context; precision is the proportion of responses assigned to that context that were correct; F1 is the harmonic mean of recall and precision. “Proportion no idea” is the proportion of sounds from each true context assigned to the “No Idea” context category.

| Context of vocalization | Number of vocalizations scored | recall | precision | F1 | Proportion no idea |
|---|---|---|---|---|---|
| Restrain | 190 | 0.226 | 0.272 | 0.247 | 0.016 |
| Castration | 152 | 0.164 | 0.301 | 0.212 | 0.044 |
| Run | 152 | 0.158 | 0.183 | 0.170 | 0.196 |
| Positive Conditioning | 161 | 0.124 | 0.128 | 0.126 | 0.139 |
| Surprise | 147 | 0.116 | 0.069 | 0.087 | 0.226 |
| Short Reunion | 156 | 0.109 | 0.092 | 0.100 | 0.188 |
| Fighting | 172 | 0.105 | 0.086 | 0.095 | 0.080 |
| Crushing | 164 | 0.098 | 0.107 | 0.102 | 0.104 |
| Enriched | 153 | 0.098 | 0.090 | 0.094 | 0.173 |
| Short Isolation | 163 | 0.080 | 0.088 | 0.084 | 0.189 |
| Before Nursing | 141 | 0.078 | 0.065 | 0.071 | 0.238 |
| After Nursing | 135 | 0.074 | 0.064 | 0.069 | 0.258 |
| Barren | 137 | 0.051 | 0.062 | 0.056 | 0.138 |
| Negative Conditioning | 151 | 0.046 | 0.054 | 0.050 | 0.161 |
| Novel Object | 176 | 0.045 | 0.053 | 0.049 | 0.107 |
| Huddling | 152 | 0.039 | 0.037 | 0.038 | 0.216 |
| Missed Nursing | 148 | 0.034 | 0.029 | 0.031 | 0.196 |
| Long Isolation | 155 | 0.026 | 0.033 | 0.029 | 0.193 |

**Table S4. Valence-level performance in the Wave 2 forced-choice valence-rating task.** Values are calculated after excluding duplicate trials. Presumed valence from Briefer et al., 2022a was binary, whereas participants rated each vocalization on a five-point ordinal scale from very negative to very positive, with an additional “No Idea Valence” option. “Precision all” is calculated separately for each non-neutral response category on the five-point scale and indicates the proportion of responses in that category that matched the corresponding presumed valence direction. For example, “Very negative” and “Negative” responses were considered correct for source-negative sounds, whereas “Positive” and “Very positive” responses were considered correct for source-positive sounds. “Precision binary” is calculated after collapsing very negative and negative responses into “negative,” and positive and very positive responses into “positive.” Recall and F1 are also calculated on this collapsed binary scale after excluding neutral and “No Idea Valence” responses. “Proportion neutral” and “Proportion no idea” indicate the proportion of sounds from each presumed valence class assigned to the neutral or “No Idea Valence” response categories, respectively.

| Valence score | N scored all | Precision all | N scored binary | Precision binary | recall | F1 | Proportion neutral | Proportion no idea |
|---|---|---|---|---|---|---|---|---|
| **Very positive** | 286 | 0.469 | | | | | | |
| **Positive** | 825 | 0.470 | 1111 | 0.470 | 0.673 | 0.553 | 0.309 | 0.136 |
| **Negative** | 726 | 0.696 | 1004 | 0.747 | 0.560 | 0.640 | 0.216 | 0.068 |
| **Very negative** | 278 | 0.881 | | | | | | |

**Table S5. Associations between conventional acoustic structure and human and neural-network similarity structure.** Pearson Mantel correlations compare context-level acoustic distance matrices with similarity structures derived from Wave 1 free classification, Wave 2 forced-choice context classification and valence rating, and context- and valence-trained neural-network representations. Acoustic distances were calculated from standardized context-level values of call duration, amplitude-modulation rate, median spectral frequency (Q50), and Wiener entropy, either jointly or separately. All tests used 9,999 permutations. False-discovery-rate (FDR) correction was applied across the four individual acoustic-feature tests within each target structure; the combined four-feature test was treated as the overall comparison and was not included in this correction.

| Target structure | Acoustic structure | Mantel correlation | FDR-adjusted P |
|---|---|---|---|
| **AI Context Model** | Combined 4 features | 0.682 | |
| | Duration | 0.490 | 0.00027 |
| | Amplitude modulation rate | 0.369 | 0.0026 |
| | Median spectral frequency | 0.514 | 2.00E-04 |
| | Wiener entropy | 0.488 | 2.00E-04 |
| **AI Valence Model** | Combined 4 features | 0.648 | |
| | Duration | 0.724 | 2.00E-04 |
| | Amplitude modulation rate | 0.097 | 0.2181 |
| | Median spectral frequency | 0.677 | 2.00E-04 |
| | Wiener entropy | 0.382 | 0.006 |
| **Human free classification** | Combined 4 features | 0.815 | |
| | Duration | 0.550 | 0.00053 |
| | Amplitude modulation rate | 0.256 | 0.0212 |
| | Median spectral frequency | 0.875 | 2.00E-04 |
| | Wiener entropy | 0.635 | 2.00E-04 |
| **Human Wave2 Context** | Combined 4 features | 0.666 | |
| | Duration | 0.629 | 2.00E-04 |
| | Amplitude modulation rate | 0.083 | 0.2592 |
| | Median spectral frequency | 0.775 | 2.00E-04 |
| | Wiener entropy | 0.497 | 0.00027 |
| **Human Wave2 Valence** | Combined 4 features | 0.750 | |
| | Duration | 0.653 | 2.00E-04 |
| | Amplitude modulation rate | 0.156 | 0.1154 |
| | Median spectral frequency | 0.837 | 2.00E-04 |
| | Wiener entropy | 0.545 | 0.00027 |

**Table S6. Acoustic predictors of human-perceived valence across sequentially adjusted cumulative-link mixed models.** Odds ratios and 95% confidence intervals represent the association between a one-standard-deviation increase in each acoustic feature and the odds of assigning a more positive valence rating. Model 1 included the four acoustic predictors; Model 2 additionally included listener characteristics; Model 3 additionally included pig characteristics; Model 4 additionally included recording characteristics; Model 5 additionally included recording context and call type. All models included random intercepts for participant and vocalization. P values were adjusted for multiple testing across the four acoustic predictors within each model using the false discovery rate (FDR) procedure.

**Abbreviations**: CI, confidence interval; FDR, false discovery rate; M1–M5, Models 1 to 5.

| Model | Acoustic feature | Odds ratio | 95% CI low | 95% CI high | FDR-adjusted P |
|---|---|---|---|---|---|
| **M1: acoustic features** | Duration | 0.425 | 0.354 | 0.510 | 6.8e-20 |
| | Amplitude modulation rate | 1.139 | 1.013 | 1.281 | 0.039 |
| | Median spectral frequency | 0.314 | 0.271 | 0.364 | 2.0e-52 |
| | Wiener entropy | 0.958 | 0.853 | 1.076 | 0.465 |
| **M2: + listener characteristics** | Duration | 0.427 | 0.356 | 0.512 | 1.3e-19 |
| | Amplitude modulation rate | 1.140 | 1.013 | 1.282 | 0.039 |
| | Median spectral frequency | 0.314 | 0.271 | 0.365 | 3.9e-52 |
| | Wiener entropy | 0.958 | 0.853 | 1.076 | 0.470 |
| **M3: + pig characteristics** | Duration | 0.464 | 0.388 | 0.554 | 4.8e-17 |
| | Amplitude modulation rate | 0.897 | 0.785 | 1.026 | 0.149 |
| | Median spectral frequency | 0.343 | 0.296 | 0.397 | 1.5e-45 |
| | Wiener entropy | 0.985 | 0.879 | 1.104 | 0.799 |
| **M4: + recording characteristics** | Duration | 0.506 | 0.427 | 0.598 | 3.4e-15 |
| | Amplitude modulation rate | 0.876 | 0.771 | 0.995 | 0.056 |
| | Median spectral frequency | 0.330 | 0.286 | 0.381 | 5.0e-51 |
| | Wiener entropy | 0.917 | 0.823 | 1.022 | 0.118 |
| **Model 5: + context / call type** | Duration | 0.589 | 0.497 | 0.698 | 2.2e-09 |
| | Amplitude modulation rate | 0.982 | 0.866 | 1.113 | 0.772 |
| | Median spectral frequency | 0.518 | 0.428 | 0.626 | 5.2e-11 |
| | Wiener entropy | 0.947 | 0.844 | 1.063 | 0.472 |

**Table S7. Context-level distribution of eligible sounds after filtering and sounds presented in Wave 1 and Wave 2.** This table reports, for each recording context, the number of eligible sounds remaining after filtering and the number of unique sounds from that context that were presented in Wave 1 and Wave 2. The table is included to assess the balance of the available and displayed stimulus pool across recording contexts.

| context | Sounds from Briefer et al. 2022b | Sounds after filtering | Sounds displayed in wave1 | Sounds displayed in wave2 |
|---|---|---|---|---|
| **After Nursing** | 79 | 56 | 56 | 53 |
| **Barren** | 585 | 491 | 300 | 145 |
| **Before Nursing** | 26 | 25 | 25 | 25 |
| **Castration** | 295 | 206 | 183 | 112 |
| **Crushing** | 146 | 125 | 122 | 95 |
| **Enriched** | 336 | 239 | 201 | 130 |
| **Fighting** | 57 | 24 | 24 | 24 |
| **Huddling** | 74 | 52 | 52 | 52 |
| **Long Isolation** | 1553 | 501 | 285 | 156 |
| **Missed Nursing** | 45 | 21 | 21 | 21 |
| **Negative Conditioning** | 119 | 4 | 4 | 4 |
| **Novel Object** | 333 | 74 | 74 | 67 |
| **Positive Conditioning** | 119 | 5 | 5 | 5 |
| **Restrain** | 927 | 379 | 264 | 144 |
| **Run** | 737 | 527 | 298 | 156 |
| **Short Isolation** | 1085 | 400 | 266 | 157 |
| **Short Reunion** | 354 | 2 | 2 | 2 |
| **Surprise** | 17 | 9 | 9 | 9 |
| **Dolphin** | 0 | 1 | 1 | 0 |

**Dataset S1 (separate file). Wave 1 free-classification responses to pig vocalizations.**

Participant- and sound-level data from the Wave 1 free-classification task. 224 Participants listened to 40 pig vocalizations without predefined response categories and sorted the sounds into self-defined groups. The dataset includes participant metadata, sound identifiers, playback information, participant-generated category labels, cleaned labels used for analysis, recording-context information, and acoustic-quality variables derived during preprocessing. These data were used to quantify spontaneous perceptual organization of pig vocalizations, participant-generated semantic labels, replay behavior, within-context consistency, and human-derived similarity matrices. The data are publicly available at https://doi.org/10.17605/OSF.IO/Q89NR (https://osf.io/q89nr/files/osfstorage).

**Dataset S2. Wave 2 forced-choice context classification and valence rating of pig vocalizations.**

Participant- and sound-level data from the Wave 2 forced-choice task. 159 Participants classified 25 pig vocalizations using predefined recording-context categories and predefined valence scores. The dataset includes participant metadata, sound identifiers, playback information, selected context and valence scores, recording-context labels, context-associated valence labels, duplicate-file indicators, and acoustic-quality variables derived during preprocessing. These data were used to assess context-classification accuracy, valence-rating accuracy, replay behavior, human context and valence similarity matrices, and comparisons with the Wave 1 free-classification results. The data are publicly available at https://doi.org/10.17605/OSF.IO/Q89NR (https://osf.io/q89nr/files/osfstorage).

**Software S1 (separate file). Analysis code for preprocessing, human-listener analyses, language analyses, machine-learning analyses, and reduced-set sensitivity analyses.**

This software file contains the documented R and Python scripts used to reproduce the analyses reported in the manuscript and Supporting Information. The scripts are numbered according to the analysis workflow: preprocessing and sound-quality filtering, Wave 1 free-classification analyses, language/semantic-label analyses, Wave 2 forced-choice context and valence analyses, machine-learning model training, embedding extraction and t-SNE/clustering analyses, and the reduced-set sensitivity analysis excluding castration, restraint, fighting, and crushing contexts. The scripts are publicly available at https://doi.org/10.17605/OSF.IO/Q89NR (https://osf.io/q89nr/files/osfstorage).

**SI References**


1. Briefer, E. F., Sypherd, C. C., Linhart, P., Leliveld, L. M. C., Padilla de la Torre, M., Read, E. R., Guérin, C., Deiss, V., Monestier, C., Rasmussen, J. H., Špinka, M., Düpjan, S., Boissy, A., Janczak, A. M., Hillmann, E., & Tallet, C. (2022a). Classification of pig calls produced from birth to slaughter according to their emotional valence and context of production. *Scientific reports*, *12*(1), 3409. https://doi.org/10.1038/s41598-022-07174-8
2. Briefer, E. F., Sypherd, C. C. R., Leliveld, L. M. C., Padilla de la Torre, M., & Tallet, C. (2022b). The Soundwel Database: a labeled pig vocalization repository [Data set]. *Scientific Reports*, *12*, 3409.
