## Supplementary material for "Pig vocalizations contain shared acoustic structure for humans and machines, but limited correspondence with presumed affective valence": Figure S18

ETH zürich

### Information and consent form

Human classification of pig vocalizations

**Contact person if you have any questions:**
Wim Gorssen, ETH Zürich

**Data Protection Officer ETH Zurich:**
Tomislav Mitar

**We would like to ask you if you are willing to participate in our research project. Your participation is voluntary. Please read the text below carefully before proceeding.**

#### What is investigated and how?

The aim of this study is to investigate how humans classify pig vocalizations. You will be asked to classify pig vocalizations and afterwards you will receive some questions.

#### Who can participate?

You may participate in this study if you are at least 18 years old and have sufficient proficiency in the language of the questionnaire to understand the instructions and questions. You should be able to listen to audio recordings using headphones or speakers in a quiet environment.

*You should not participate if you have a hearing impairment that would substantially affect your ability to perceive or distinguish sounds.*

#### What am I supposed to do as a participant?

If you agree to participate, you will be asked to listen to a series of recorded pig vocalizations and to classify them. You will also be asked to answer short questions about your background, experience with pigs, and basic demographic information. The study is conducted online and takes approximately 20-30 minutes to complete.

#### What are my rights during participation?

Your participation in this study is voluntary. You can withdraw your participation at any time without giving reasons and without any disadvantage.

#### What risks and benefits can I expect?

There are no known risks associated with participation. Some participants may find listening to animal sounds mildly uncomfortable. You may pause or stop the study at any time.

#### Will I be compensated for my participation?

You will not receive financial compensation for participating in this study. Participation is entirely voluntary.

#### What data is collected from me and how is it used?

No identifying personal data are collected in this study. This means that your name, email address, IP address, or any other information that could directly identify you is not recorded.

The following research data are collected:

- Responses to questionnaire items (e.g. background, experience with animals, pet ownership, and demographic categories)
- Responses to the vocalization classification tasks

No audio, video, or voice recordings of participants are collected. The results of the study will be published in scientific journals and presented at academic conferences. Only aggregated results will be reported, and no conclusions about individual participants can be drawn from any publication.

#### Who is funding the study?

This study has no external funding. It is conducted as part of academic research at ETH Zurich without dedicated financial support.

#### Who reviewed the study?

This study was reviewed and approved by the ETH Zurich Ethics Committee under application number 26 ETHICS-004.

#### Complaints office

The secretariat of the ETH Zurich Ethics Committee is available to help you with complaints in connection with your participation in the study. Contact: or 0041 44 632 85 72.

#### Electronic consent form

**By clicking the box below, I, the participant, confirm that:**

- I have read and understood the study information.
- I had enough time to decide about my participation and am taking part in the study voluntarily.
- I fulfil the stated conditions for participation.
- I consent that the data described above may be collected from me and used as described.
- I know that the data collected from me may be published in anonymised form.
- I know that I can stop my participation any time.

☐ I agree to participate in this study.

Continue

Project Oink

GB English

### Project Oink

Classify each pig vocalization into categories that make sense to you. You can click on the sounds to hear them under 'Available Audio Files'. You can create categories under 'Create New Box'. You can drag the audio files in your created categories.

#### Create New Box

Enter box name...

Create Box

#### Your Boxes

No boxes yet. Create your first box to get started!

#### Available Audio Files

#0 #1 #2
#3 #4 #5
#6 #7 #8
#9 #10 #11
#12 #13 #14
#15 #16 #17
#18 #19 #20
#21 #22 #23
#24 #25 #26
#27 #28 #29
#30 #31 #32
#33 #34 #35

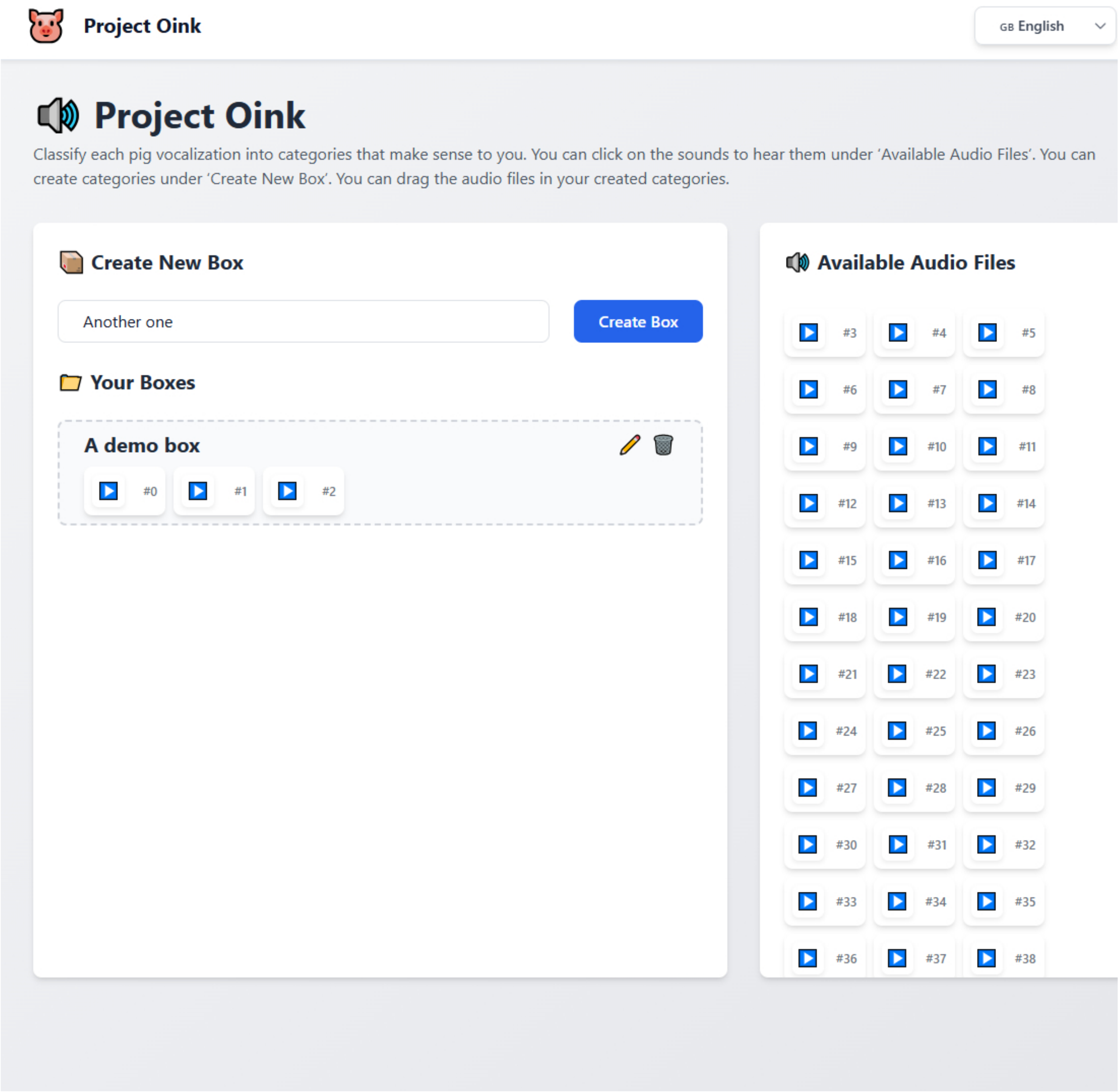
Project Oink
GB English
Project Oink
Classify each pig vocalization into categories that make sense to you. You can click on the sounds to hear them under 'Available Audio Files'. You can create categories under 'Create New Box'. You can drag the audio files in your created categories.
Create New Box
Another one
Create Box
Your Boxes
A demo box
#0
#1
#2
Available Audio Files
#3
#4
#5
#6
#7
#8
#9
#10
#11
#12
#13
#14
#15
#16
#17
#18
#19
#20
#21
#22
#23
#24
#25
#26
#27
#28
#29
#30
#31
#32
#33
#34
#35
#36
#37
#38

Project Oink

#### Survey Questions

**What is your primary professional background? (Select 1-3)**

- [ ] Pig farmer
- [ ] Animal caretaker
- [ ] Veterinarian
- [ ] Animal scientist or ethologist
- [ ] Animal welfare or rights professional
- [ ] Bioacoustician
- [ ] Musician
- [ ] No relevant professional background
- [ ] Other, please specify

- [ ] Prefer not to say

**In the past 12 months, how often have you had direct contact with pigs?**

- ○ Daily
- ○ Weekly
- ○ Monthly
- ○ Yearly
- ○ No contact the last year
- ◉ Prefer not to say

**How would you rate your level of experience with pigs? (0-10)**

0-10 ☐ Prefer not to say

**How would you rate your level of experience with sound analysis or auditory discrimination?**

- ○ None
- ○ Basic
- ○ Intermediate
- ○ Advanced
- ○ Expert
- ◉ Prefer not to say

GB English

**What is your age group?**

- ○ 18-24
- ○ 25-34
- ○ 35-44
- ○ 45-54
- ○ 55-64
- ○ 65-74
- ○ 75-84
- ○ 85+
- ◉ Prefer not to say

**What is your gender?**

- ○ Female
- ○ Male
- ○ Other
- ◉ Prefer not to say

**Do you currently own or regularly care for a pet animal?**

- ○ Yes
- ○ No
- ◉ Prefer not to say

Submit

Project Oink

GB English

### Part 2 – Context of pig sounds

Assign each audio file to the box that best matches how you perceive the sound.

#### Context Definitions (Click to Expand/Collapse)

–

In the next part, you will be asked to classify 25 pig vocalizations into the context in which you think they are produced. In total, there are 18 different contexts from which pig vocalizations where recorded, with following definition:

**After Nursing:** Approach to the sow's head after completing nursing

**Before Nursing:** Approach to the sow's head before nursing

**Crushing:** A pig lying on its side while a human applies downward pressure to press the body against the floor
**Fighting:** Competition among pigs
**Long Isolation:** Social isolation in a familiar arena for 8 minutes
**Negative Conditioning:** Exposure to a negatively conditioned familiar arena without another pig
**Positive Conditioning:** Exposure to a familiar, positively conditioned arena without another pig
**Reunion:** Reuniting with other pigs after isolation

**Short Isolation:** Social isolation in a familiar arena for 3 minutes

**Barren environment:** Exposure to a familiar, non-enriched arena together with another pig
**Castration:** Surgical removal of the testes without anaesthesia, performed by a human while the animal is restrained
**Enriched environment:** Exposure to a familiar, enriched arena together with another pig
**Huddling:** Physical grouping in close contact with other pigs
**Missed Nursing:** Struggling to reach a teat for nursing
**Novel Object:** Exposition alone to a novel object in a novel arena

**Restrain:** physical restraint by a human, either by holding the animal in the arms or by turning the animal onto its back and holding the forelegs and abdomen
**Run:** Running from the pen to a familiar arena or vice versa together with another pig
**Surprise:** Being surprised by the arrival of a person

#### Boxes

##### Run
#0

##### Novel Object
#1

##### Before Nursing
#5

##### Huddling
#2

##### Short Isolation
#3

##### Barren environment
#4

##### Positive Conditioning
#15

##### Castration
#18

##### Missed Nursing
Drop audio files here

##### Negative Conditioning
#12

#### Available Audio Files

#6 #7 #8

#9 #10 #11

#13 #14 #16

#20 #23 #24

Project Oink

#### Survey Questions

What is your primary professional background? (Select 1-3)

- [ ] Pig farmer
- [ ] Animal caretaker
- [ ] Veterinarian
- [ ] Animal scientist or ethologist
- [ ] Animal welfare or rights professional
- [ ] Bioacoustician
- [ ] Musician
- [ ] No relevant professional background
- [ ] Other, please specify

- [ ] Prefer not to say

GB English

In the past 12 months, how often have you had direct contact with pigs?

- ( ) Daily
- ( ) Weekly
- ( ) Monthly
- ( ) Yearly
- ( ) No contact the last year
- (•) Prefer not to say

How would you rate your level of experience with pigs? (0-10)

0-10 [ ] Prefer not to say

How would you rate your level of experience with sound analysis or auditory discrimination?

- ( ) None
- ( ) Basic
- ( ) Intermediate
- ( ) Advanced
- ( ) Expert
- (•) Prefer not to say

What is your age group?

- ( ) 18-24
- ( ) 25-34
- ( ) 35-44
- ( ) 45-54
- ( ) 55-64
- ( ) 65-74
- ( ) 75-84
- ( ) 85+
- (•) Prefer not to say

What is your gender?

- ( ) Female
- ( ) Male
- ( ) Other
- (•) Prefer not to say

Do you currently own or regularly care for a pet animal?

- ( ) Yes
- ( ) No
- (•) Prefer not to say

Submit

Project Oink

GB English

### Part 1 - How positive or negative does this pig sound?

Assign each audio file to the box that best matches how you perceive the sound.

#### Boxes

**Very negative**
#0

**Negative**
#1

**Neutral**
#2

**Positive**
#3

**Very positive**
#4

**No idea**
#5 #6

#### Available Audio Files

#7 #8 #9
#10 #11 #12
#13 #14 #15
#16 #17 #18
#19 #20 #21
#22 #23 #24

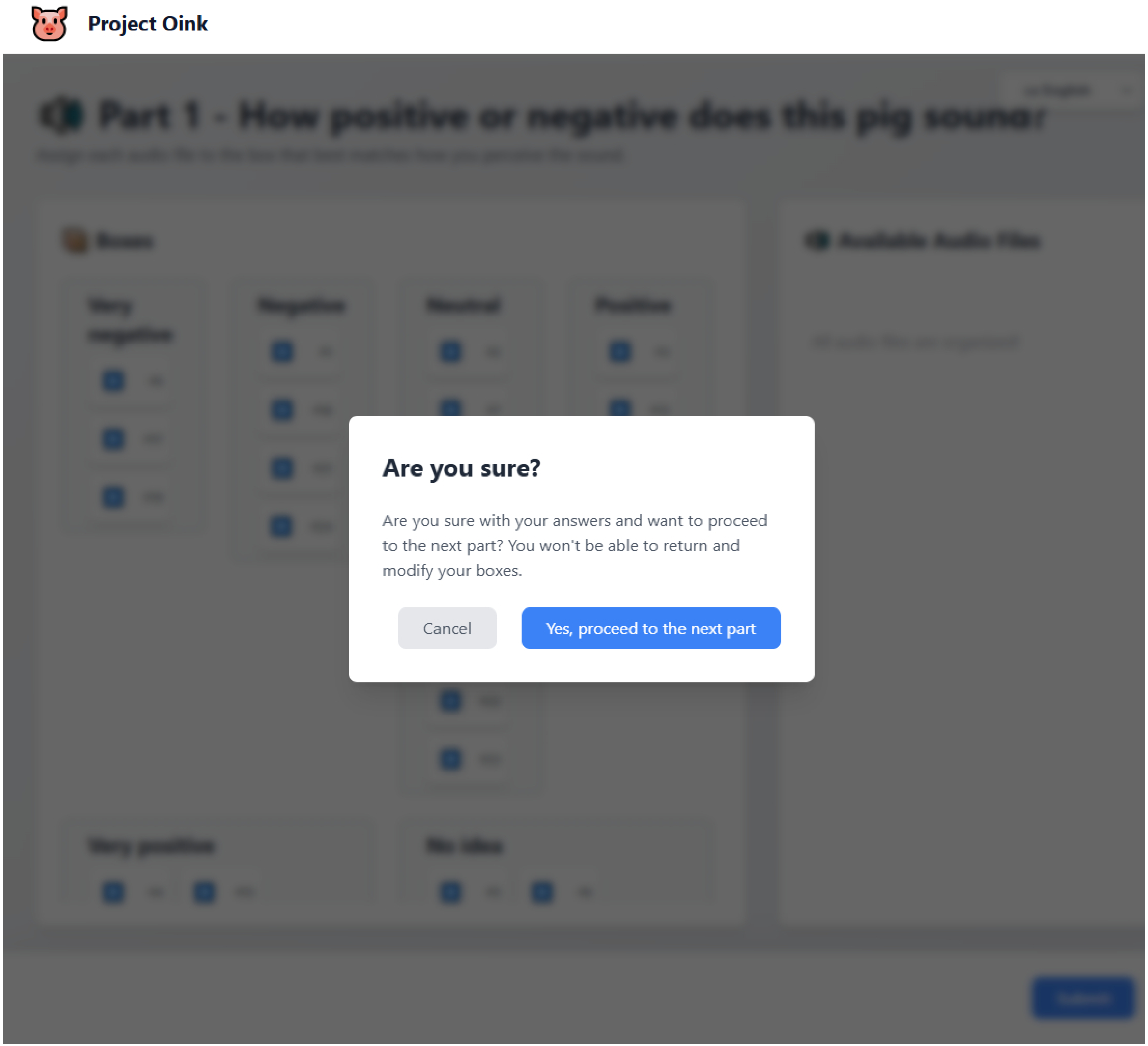
Project Oink
Are you sure?
Are you sure with your answers and want to proceed to the next part? You won't be able to return and modify your boxes.
Cancel
Yes, proceed to the next part

Project Oink

GB English

### Thank You!

Your session has been completed. You can now close this page.

Your responses have been submitted successfully. We appreciate your participation in this study.
